# Evolution of SARS-CoV-2 T cell responses as a function of multiple COVID-19 boosters

**DOI:** 10.1101/2025.01.08.631842

**Authors:** Ricardo da Silva Antunes, Vicente Fajardo-Rosas, Esther Dawen Yu, Rosa Isela Gálvez, Adam Abawi, E. Alexandar Escarrega, Amparo Martínez-Pérez, Emil Johansson, Benjamin Goodwin, April Frazier, Jennifer M. Dan, Shane Crotty, Grégory Seumois, Daniela Weiskopf, Pandurangan Vijayanand, Alessandro Sette

## Abstract

The long-term effects of repeated COVID-19 vaccinations on adaptive immunity remain incompletely understood. Here, we conducted a comprehensive three-year longitudinal study examining T cell and antibody responses in 78 vaccinated individuals without reported symptomatic infections. We observed distinct dynamics in Spike-specific humoral and cellular immune responses across multiple vaccine doses. While antibody titers incrementally increased and stabilized with each booster, T cell responses rapidly plateaued, maintaining remarkable stability across CD4+ and CD8+ subsets. Notably, approximately 30% of participants showed CD4+ T cell reactivity to non-Spike antigens, consistent with asymptomatic infections. Single-cell RNA sequencing revealed a diverse landscape of Spike-specific T cell phenotypes, with no evidence of increased exhaustion or significant functional impairment. However, qualitative changes were observed in individuals with evidence of asymptomatic infection, exhibiting unique immunological characteristics, including increased frequencies of Th17-like CD4+ T cells and GZMKhi/IFNR CD8+ T cell subsets. Remarkably, repeated vaccinations in this group were associated with a progressive increase in regulatory T cells, potentially indicating a balanced immune response that may mitigate immunopathology. By regularly stimulating T cell memory, boosters contribute to a stable and enhanced immune response, which may provide better protection against symptomatic infections.

## INTRODUCTION

Vaccines against SARS-CoV-2 played a critical role in efforts to combat the COVID-19 pandemic (Andrews et al., 2022b; Baden et al., 2021; Polack et al., 2020). In addition to the primary series of SARS-CoV-2 vaccinations that became available in the US at the end of 2020 and the beginning of 2021, additional booster vaccinations have been recommended and widely utilized, with the goal of enhancing and prolonging protection against infection and disease, particularly in response to waning immunity over time, and the emergence of SARS-CoV-2 variants (Andrews et al., 2022a; Atmar et al., 2022; Chemaitelly et al., 2023).

Primary vaccination is in general associated with high levels of neutralizing antibodies whose levels drop significantly within months (Sette and Crotty, 2022; Srivastava et al., 2024; Zhang et al., 2022a). In contrast, T cell responses observed after vaccination are more sustained and persist for at least six months (Moss, 2022; Sette et al., 2023; Zhang *et al*., 2022a). However, the long-term impact of multiple booster vaccinations on the persistence of adaptive immune responses, is poorly understood. At the level of antibody response, multiple boosters affect both magnitude, quality, and kinetics of decay of antibody titers (Srivastava *et al*., 2024).

Both CD4+ helper and CD8+ cytotoxic T cells play an important role in protection against hospitalization, severe disease, and death (Bertoletti et al., 2022; Kedzierska and Thomas, 2022; Moss, 2022). CD4+ and CD8+ T cell responses correlate with more rapid viral clearance (Grifoni et al., 2020b; Tan et al., 2021), and early responses are associated with milder disease outcomes (Tan *et al*., 2021; Tarke et al., 2022). T cell responses are largely preserved and effectively cross-recognize variants of concern (VoC), in contrast to neutralizing antibody responses, which are substantially impacted in the context of emerging variants (Geers et al., 2021; Riou et al., 2022; Tarke et al., 2021). Akin to T cells generated during acute SARS-CoV-2 infection, COVID-19 vaccine responses persist over extended periods of time (Dan et al., 2021a; Moss, 2022; Zhang *et al*., 2022a). However, little is known about the evolution of memory phenotypes and functionality over time, particularly in the context of repeated booster vaccinations.

While studies have examined T cell response durability after booster vaccinations (Arunachalam et al., 2023; Reinscheid et al., 2022), it remains unclear whether booster vaccinations increase the magnitude of long-term T cell memory, or if a plateau level no longer impacted by further boosters is reached. Booster shots may also be associated with modulation of the quality of T cell responses. Specifically, T cell responses can become more robust and diverse with repeated antigen exposure (Cai et al., 2023; Painter et al., 2023; Tarke et al., 2024), but concerns have also been raised regarding the possibility that repeated vaccinations might be associated with exhaustion and progressive lack of functionality (Azim Majumder and Razzaque, 2022; Boretti, 2024; Jo et al., 2023). Additionally, the impact of previous asymptomatic infections, which are becoming increasingly common even in fully vaccinated populations, on both the magnitude and durability of these responses, remains unknown (Le Bert and Samandari, 2024; Shaikh et al., 2023).

Our study, conducted over a 3-year period (Jan 2021-Jan 2024) during the COVID-19 pandemic period, addresses these critical questions by examining the evolution of Spike-specific T cell and antibody responses in a cohort of 78 vaccinated individuals who did not report symptomatic infections and did not test positive throughout the study period.

## RESULTS

### Vaccination Cohort Characteristics

We enrolled an observational cohort of several hundred participants followed over a 3-year period (Jan 2021-Jan 2024). The participants were vaccinated with mRNA vaccines, either mRNA-1273 (Moderna), BNT162b2 (Pfizer/BioNTech) or a combination of both, or NVX-CoV2373 (Novavax). Herein, we characterize the evolution of SARS-CoV-2-specific adaptive immune responses as a function of the number of vaccination doses in a subset of 78 participants who did not report COVID-19-related symptoms nor tested positive for SARS-CoV-2 at any timepoint (**Fig 1A**). The majority of participants (77/78) received either Pfizer or Moderna COVID-19 vaccines, while one participant received Novavax. All participants received up to four different doses. The overall characteristics of the cohort are summarized in **Table S1** and detailed in the STAR Methods section. The cohort was further divided into 3 different groups as a function of the number of vaccination doses, i.e., full vaccination or Dose 2 (D2), first booster or Dose 3 (D3) and second booster or Dose 4 (D4) (**Fig 1B**). All donors (100%) received at least 2 vaccinations, while 88.4% and 74.3% received 3 and 4 vaccine doses, respectively.

**Figure 1.**
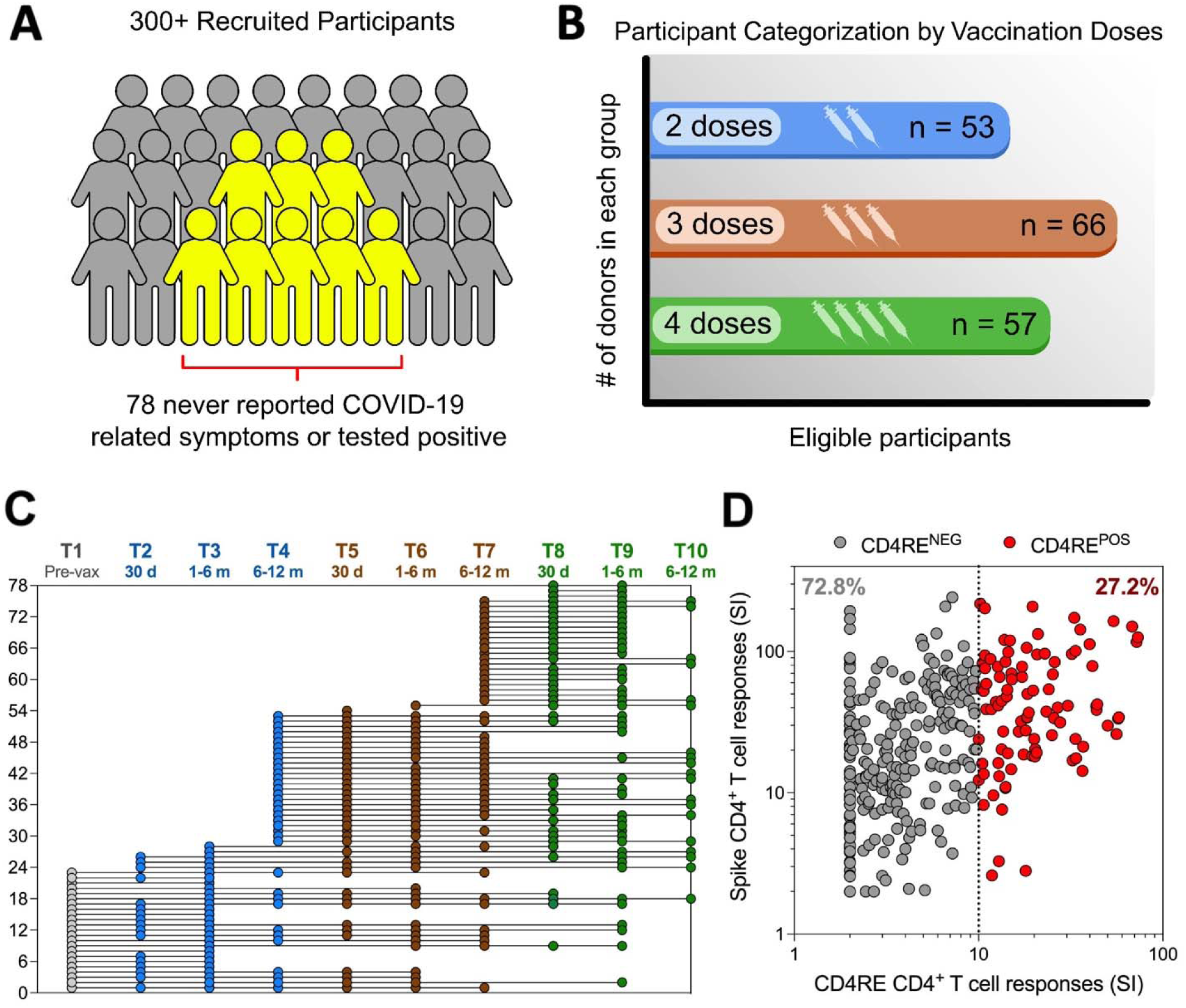
Study design and prevalence of asymptomatic infections. (**A**) Overall cohort surveillance for selection of 78 participants without COVID-19 symptoms or previous evidence of infection. (**B**) Participants categorization by the number of COVID- 19 vaccine doses administered (color-coded). The number of donors included in each group is indicated and the groups are not mutually exclusive, allowing individual donors to be represented in multiple groups. (**C**) Multiple blood donations were collected from each subject at various timepoints (T) post-vaccination. The y axis represents each individual participant, and the x axis represents different time points at which their blood was collected in a longitudinal follow-up throughout the study timeline (Jan 2021-Jan 2024). Samples within the period of each vaccine dose are color-coded. Sampling timepoints include pre-vaccination (T1), 30 days after full vaccination (Dose 2) (T2), 30 days after the first booster (Dose 3) (T5), and 30 days after the second booster (Dose 4) (T8). Time intervals between vaccinations encompass T3, T6 and T9 (1-6 months after Dose 2, 3 and 4, respectively) and T4, T7 and T10 (6-12 months after Dose 2, 3 and 4, respectively). (**D**) CD4+ T cell responses to Spike and CD4RE MPs were measured as percentage of AIM+ (OX40+CD137+) CD4+ T cells and plotted in 2 dimensions as stimulation index (SI). Each dot represents a sample of a donor for a given timepoint. Percentages denote the frequency of samples that are either negative (CD4RE^NEG^) or positive (CD4RE^POS^) for CD4RE reactivity using a cut-off of SI=10 (dotted line).

Samples associated with each visit were organized according to the temporal window of collection or timepoints (T): pre-vaccination (T1), and around 30 days after D2, D3, and D4 (T2, T5, and T8, respectively). The intervals between vaccinations were further segmented into periods of 1-6 months (T3, T6, and T9) or 6-12 months (T4, T7, and T10) post-D2, D3, and D4 vaccinations. **Fig 1C** provides a schematic overview depicting the individual donors analyzed, and their relationship to the number of immunizations and the time of sample collection within the study cohort (see STAR Methods for more details). The breakdown of assays performed per donor is detailed in **Table S2**. In summary, this study consists of three participant groups, each of which received a different number of COVID-19 vaccinations: D2, D3, and D4, with 53, 66, and 57 donors respectively in each group; Of these, 33 donors have longitudinal visits across all three groups.

### Stratification based on T cell reactivity to non-S antigens

We previously developed an immunodiagnostic tool to identify prior exposure to SARS-CoV-2 (Yu et al., 2022b), by measuring CD4+ T cell reactivity to two peptide megapools (MP): one covering the entire sequence of the ancestral Spike protein and another consisting of experimentally validated CD4 epitopes from the remainder of the SARS-CoV-2 proteome, known as CD4RE (non-Spike epitopes). Here, we applied this methodology to identify vaccinated individuals who likely experienced SARS-CoV-2 infection, based on detectable CD4RE reactivity. Indeed, even though all samples were obtained from donors who did not self-report SARS-CoV-2 infections or COVID-19- related symptoms at any time point associated with each blood donation, an activation-induced cell marker (AIM) assay with a very stringent stimulation index (SI) cutoff (10- fold) detected CD4RE reactivity in 27.2 % of the total samples (**Fig 1D**; hereafter CD4RE^POS^ samples).

In summary, these data show that a large fraction of samples from vaccinated individuals in the cohort, who did not report symptomatic infections or positive SARS-CoV-2 test results, nevertheless had detectable reactivity to CD4RE, suggesting these subjects experienced asymptomatic infections. These findings are consistent with the high incidence of asymptomatic infections reported in meta-analyses, serological studies, and T cell response studies conducted during the Delta and Omicron waves (Tarke *et al*., 2024; Wang et al., 2023; Yu et al., 2022c). To address this confounding factor, the three cohorts were further analyzed taking sample CD4RE reactivity into consideration.

### Evolution of antibody responses over repeated exposures and vaccinations

SARS-CoV-2 Spike RBD antibodies titers over the course of the 3 years were determined as previously described (Zhang *et al*., 2022a) (**Fig 2**), in a subset of 68 donors (**Table S2**). Seroconversion was observed in all donors after the first 2 immunizations (T2). Increase of antibody titers was also observed in all donors after both the third (T5) and fourth (T8) doses (**Fig 2A**), although subsequent doses of COVID-19 vaccination resulted in progressive smaller enhancement of the RBD titers. The fold change (FC) in titers between each immunization decreased from 588x after D2, to 10.7x after D3, and finally to 2.7x following D4 (**Fig 2B**, top panel). This pattern indicates a reduced boosting effect upon additional vaccinations.

**Figure 2.**
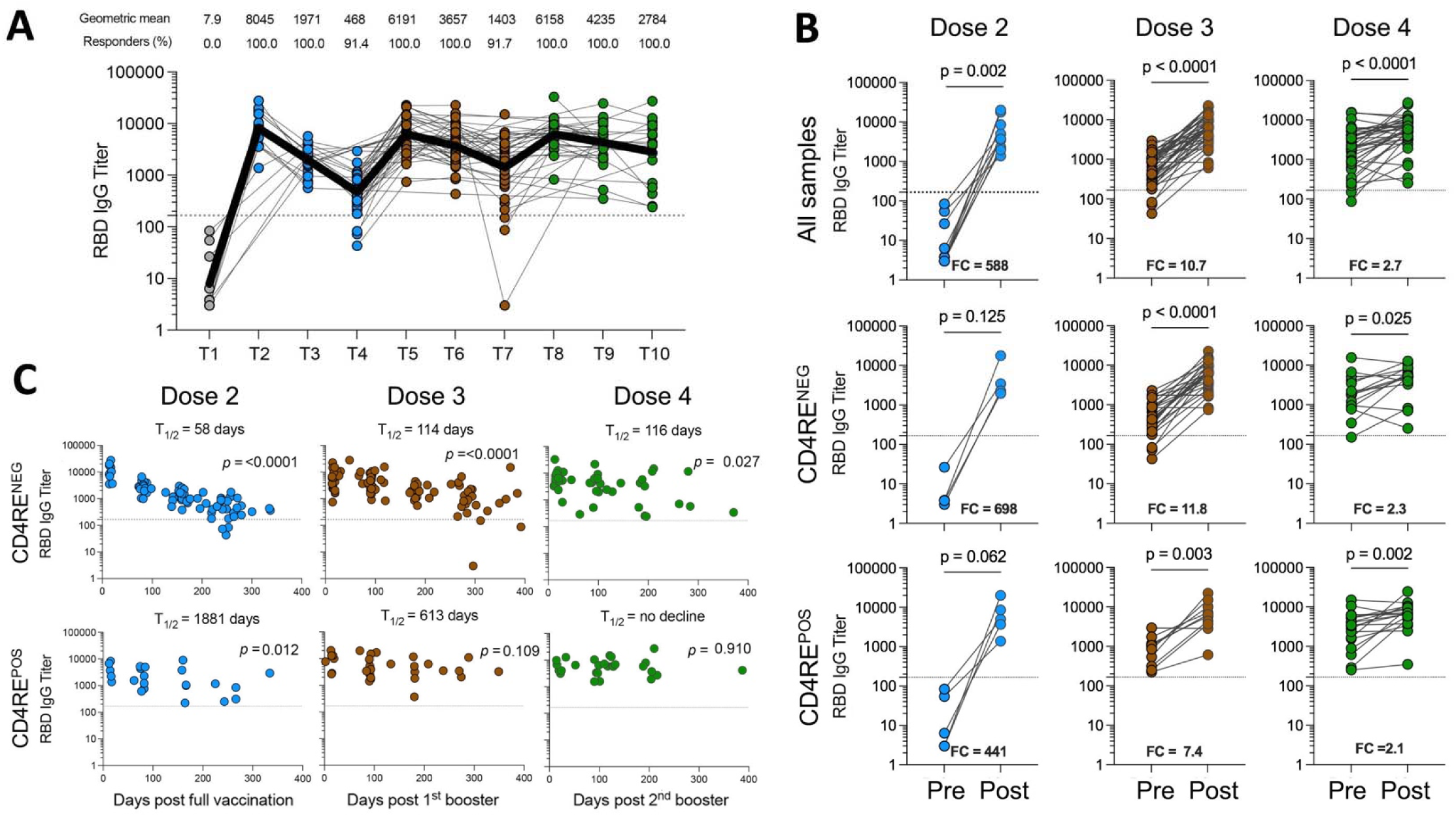
Evolution of antibody responses over repeated vaccinations and exposure. (**A**) Longitudinal monitoring of RDB IgG levels across multiple vaccinations and timepoints. Each donor is represented by a colored dot and longitudinal samples for each individual donor connected by a gray line. Black bold line represents the geometric mean. Geometric mean value and % of donors with positive response for each timepoint is indicated. (**B**) Representation of paired RDB IgG levels before and after vaccination for each dose cohort for all samples (upper panel), or CD4RE^NEG^ (middle panel) and CD4RE^POS^ (lower panel) donors. Fold change (FC) is indicated in each graphic. (**C**) Cross-sectional RBD IgG responses from time of vaccination (days) for the different dose cohorts in donors associated with CD4RE^NEG^ (upper panel) or CD4RE^POS^ (lower panel) reactivity. t_1/2_ is shown as the median half-life calculated based on linear mixed effects model. (**A-C**) Dotted lines indicate threshold of positivity. Data were analyzed for statistical significance using the paired Wilcoxon’s (**B**) or Spearman’s (**C**) test. p values are shown.

The effect of repeated vaccinations was further analyzed by plotting the available longitudinal sample pairs that had been consistently (before and after each immunization) associated with negative reactivity for CD4RE, versus having tested positive for CD4RE reactivity at any point in time before the immunization (**Fig 2B**, middle and lower panels). For individuals with presumed asymptomatic infections (i.e., CD4RE^POS^), the titer increases after the first two immunizations were considerably lower compared with CD4RE^NEG^ donors (FC=441 vs. 698), consistent with pre-existing SARS-CoV-2 Spike-specific responses. Although more modest, this difference was maintained after both the third (FC=7.4 vs. 11.8) and fourth (FC=2.3 vs. 2.1) immunizations.

We further analyzed the RBD IgG titer durability across the three cohorts over time by estimating half-lives using a previously described methodology (Yu et al., 2022a) (**Fig 2C**). As previously reported in numerous studies (Levin et al., 2021; Sette and Crotty, 2022; Srivastava *et al*., 2024; Zhang *et al*., 2022a), antibody titters declined relatively rapidly following the completion of the first 2 vaccinations. Strikingly, subsequent boosters increased durability of the antibody responses in the cohort associated with CD4RE^NEG^ responses, and following D2, antibody titers declined faster in the CD4RE^NEG^ compared to the CD4RE^POS^ cohort (58 vs. 1,881 days, respectively). The CD4RE^POS^ asymptomatic infection cohort was associated with relatively stable antibody titers (**Fig 2C**).

In summary, repeated boosters increased antibody responses both in magnitude and durability, with the additional exposure(s) associated with asymptomatic infections also contributing to more stable responses. These results also emphasize that accurate determination of a donor’s infection history and categorization accounting for asymptomatic infections, are crucial to correctly assess the impact of booster vaccinations on virus-specific antibody titers.

### Evolution of memory Spike-specific CD4^+^ and CD8+ T cell responses over repeated vaccinations and exposures

We next examined how the number of vaccinations and previous asymptomatic infections influence the magnitude and durability of CD4+ and CD8+ T cell responses, using the same subset of 68 donors previously assessed for antibody titers (**Table S2**). Specifically, we utilized the AIM assay (OX40+4-1BB+ and CD69+4-1BB+ marker combination for CD4+ and CD8+ T cells, respectively) (da Silva Antunes et al., 2023; Dan et al., 2016), to measure Spike-specific responses, using a MP of overlapping peptides covering the entire length of the Spike ancestral sequence used in Pfizer and Moderna SARS-CoV-2 vaccines (Grifoni *et al*., 2020b).

The evolution of CD4+ T cell responses was monitored over the course of 3 years (**Fig 3**). The magnitude of CD4+ T cell responses increased after the first two vaccinations and remained relatively stable throughout the course of multiple vaccinations (**Fig3A**). All vaccinees (100%) exhibited detectable CD4+ T cell responses following the initial two-dose immunization (T2), as well as after the third (T5) and fourth (T8) doses. Following the third and fourth doses of COVID-19 vaccination (**Fig 3B**, top panel), T cell responses had reached or were approaching a plateau, with high magnitude and frequency of responses still observed 6-12 months after the second (T4) and third (T7) doses, respectively. We further analyzed the effect of booster immunizations by comparing longitudinal samples with consistent CD4RE reactivity across immunizations (**Fig 3B**, middle and lower panels). CDRE^POS^ individuals showed a smaller increase after the first two immunizations compared to CD4RE^NEG^ donors (fold change of 6.8 vs. 10.7), but no impact of asymptomatic infection was observed after third and fourth doses (fold changes of 1.0 vs. 1.2 and 1.2 vs. 0.9, respectively).

**Figure 3.**
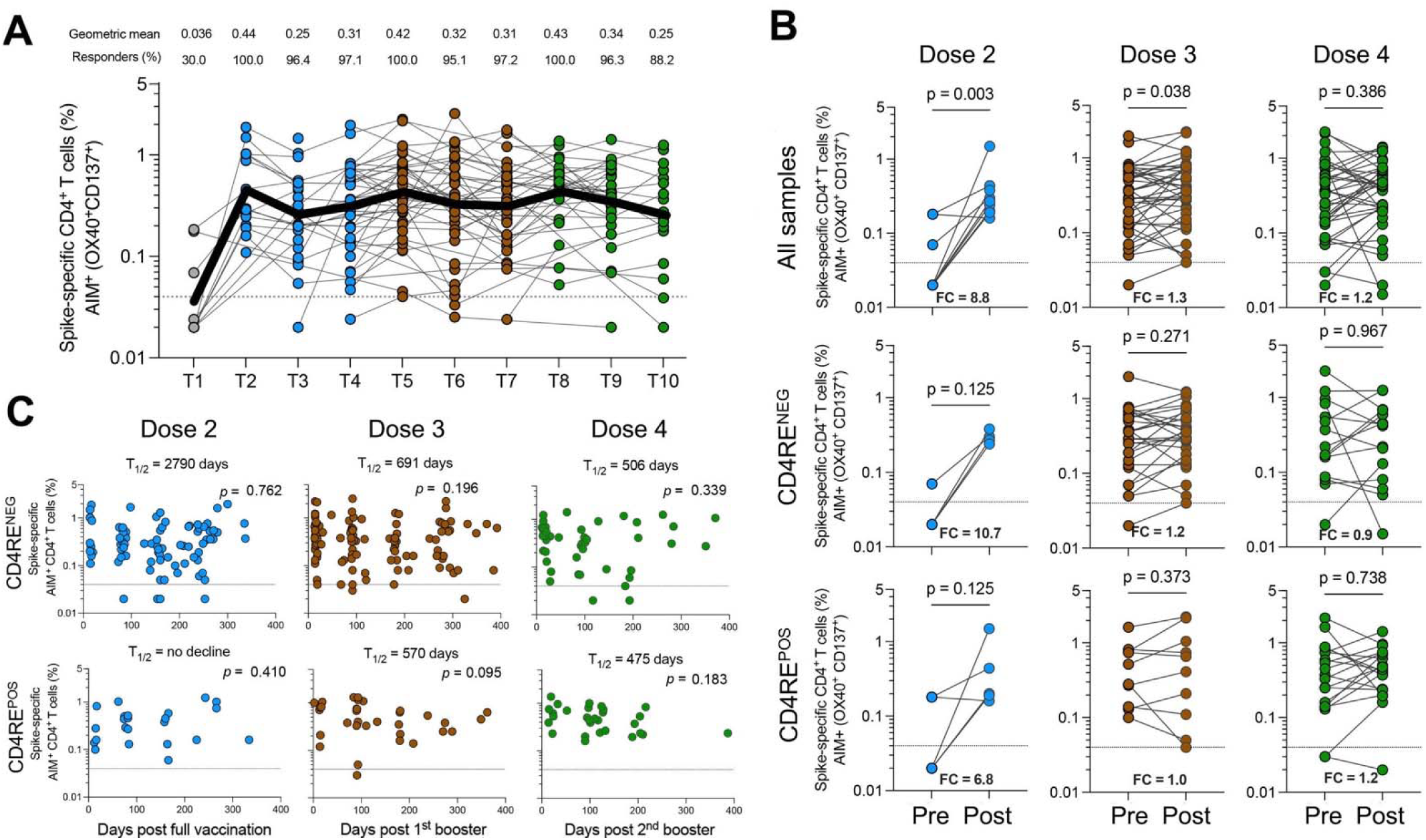
Evolution of memory Spike-specific CD4+ T cell responses over repeated exposures and vaccinations. (**A**) Longitudinal monitoring of AIM+ (OX40+CD137+) CD4+ T cell responses across multiple vaccinations and timepoints. Each donor is represented by a colored dot and longitudinal samples for each individual donor connected by a gray line. Black bold line represents the geometric mean. Geometric mean value and % of donors with positive response for each timepoint is indicated. (**B**) Representation of paired AIM+ (OX40+CD137+) CD4+ T cell responses before and after vaccination for each dose cohort for all samples (upper panel), or CD4RE^NEG^ (middle panel) and CD4RE^POS^ (lower panel) donors. Fold change (FC) is indicated in each graphic. (**C**) Cross-sectional AIM+ (OX40+CD137+) CD4+ T cell responses from time of vaccination (days) for the different dose cohorts in donors associated with CD4RE^NEG^ (upper panel) or CD4RE^POS^ (lower panel) reactivity. t_1/2_ is shown as the median half-life calculated based on linear mixed effects model. (**A-C**) Dotted lines indicate threshold of positivity. Data were analyzed for statistical significance using the paired Wilcoxon’s (**B**) or Spearman’s (**C**) test. p values are shown.

The durability of Spike-specific CD4+ T cells in each of three cohorts was also investigated, with the same methodology described for antibody responses above (**Fig 3C**). CD4^+^ T cell responses were relatively stable over the period considered. More specifically, the estimated half-lives after two, three, and four immunizations were respectively 2,790, 691, and 506 days in the CD4RE^NEG^ samples and no decline, 570, and 475 days for the CD4RE^POS^ samples. Overall, the data indicate that CD4+ T cell responses were substantially stable, irrespective of the CD4RE status.

CD8+ T cell responses, like their CD4+ counterparts, were generally stable across multiple vaccinations (**Fig 4**). Although less frequently detectable than CD4+ T cell responses, CD8+ T cell responses were nevertheless detected in the majority of donors following successive immunizations (D2=81.3%, D3=76.3%, and D4=83.3%) (**Fig 4A**). Similarly, the enhancement of CD8+ T cell immune responses following additional vaccinations was limited after the third and fourth COVID-19 vaccine doses (**Fig 4B**, top panel), and slightly more pronounced in the group of CD4RE^POS^ samples (**Fig 4B**, middle and lower panels). Like CD4+ T cell responses, Spike-specific CD8+ T cell responses were fairly stable throughout the follow up period regardless of previous asymptomatic infections (**Fig 4C**). In conclusion, T cell responses plateau more quickly than antibody responses following repeated vaccinations and asymptomatic infections and remained largely stable throughout the study follow-up period.

**Figure 4.**
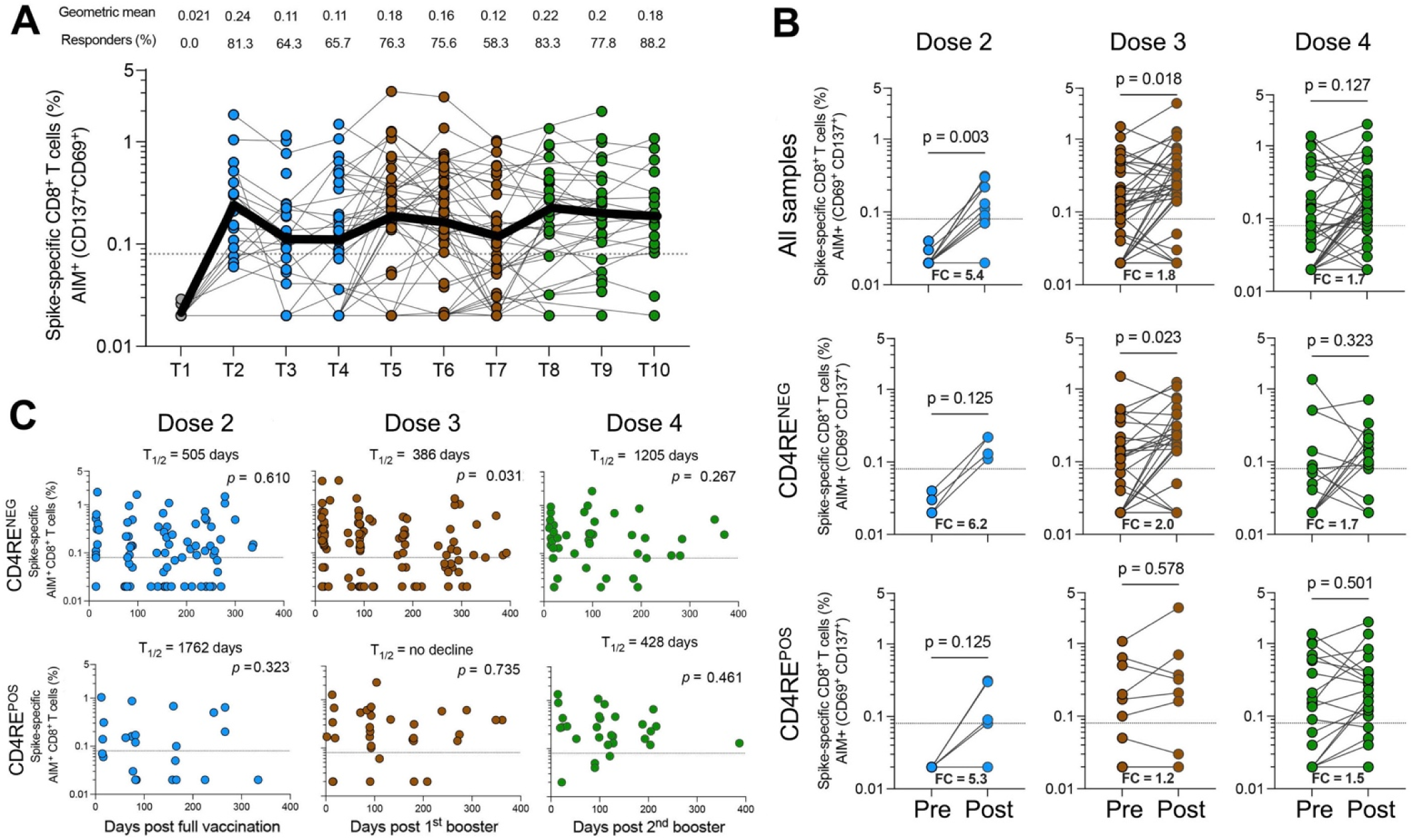
Evolution of memory Spike-specific CD8+ T cell responses over repeated exposures and vaccinations. (**A**) Longitudinal monitoring of AIM+ (CD69+CD137+) CD8+ T cell responses across multiple vaccinations and timepoints. Each donor is represented by a colored dot and longitudinal samples for each individual donor connected by a gray line. Black bold line represents the geometric mean. Geometric mean value and % of donors with positive response for each timepoint is indicated. (**B**) Representation of paired AIM+ (CD69+CD137+) CD8+ T cell responses before and after vaccination for each dose cohort for all samples (upper panel), or CD4RE^NEG^ (middle panel) and CD4RE^POS^ (lower panel) donors. Fold change (FC) is indicated in each graphic. (**C**) Cross-sectional AIM+ (CD69+CD137+) CD8+ T cell responses from time of vaccination (days) for the different dose cohorts in donors associated with CD4RE^NEG^ (upper panel) and CD4RE^POS^ (lower panel) reactivity. t_1/2_ is shown as the median half-life calculated based on linear mixed effects model. (**A-C**) Dotted lines indicate threshold of positivity. Data were analyzed for statistical significance using the paired Wilcoxon’s (**B**) or Spearman’s (**C**) test. p values are shown.

### scRNA sequencing studies to determine the evolution of memory T cell phenotypes over repeated vaccinations

The data above underline how T cell responses plateau relatively rapidly and are remarkably stable following multiple exposures. Next, we examined the effect of multiple COVID-19 vaccinations and asymptomatic infections on the quality and dynamics of Spike-specific T cell responses. Specifically, we examined the transcriptional properties of T cells at single-cell resolution in a subset of 54 donors for whom sufficient samples were available. This cohort largely overlapped with the previous one, described above, with 74.4% of the samples being shared (**Table S2**). Like in the previous experimental design, peripheral blood mononuclear cells (PBMCs) were thawed and stimulated for 24 hours with a pool of overlapping peptides spanning the entire sequence of the Spike antigen (see STAR Methods). Spike-specific CD4+ and CD8+ AIM+ T cells (OX40+4- 1BB+ and 4-1BB+CD69+, respectively) were sorted and profiled for their transcriptomes at single-cell resolution.

Our analysis resulted in a total of 1,069,657 high-quality single-cell transcriptomes from different doses and multiple timepoints. For Spike-specific CD4+ T cells, we analyzed 59, 108, and 106 distinct samples after 2, 3, and 4 vaccine doses respectively. For Spike-specific CD8+ T cells, we analyzed 65, 99, and 95 distinct samples after 2, 3, and 4 doses, respectively. The median number of high-quality cells per sample was 1,754 and 1,407 for Spike-specific CD4+ and CD8+ T cells respectively, and samples with less than 100 cells were excluded from the subsequent analyses. Overall, abundant Spike-specific T cells were recovered across multiple samples and well-balanced among different vaccine doses and timepoints (**Table S3**).

### High resolution analysis of Spike-specific CD4+ and CD8+ T cell subsets

Unbiased clustering analysis of the scRNA-seq data revealed several different clusters of Spike-specific T cells (**Fig 5**), which were annotated on the basis of their differentially expressed genes (**Table 1** and **Fig S1**). In the case of Spike-specific CD4+ T cells (**Fig 5A**), eight different clusters were identified, with the regulatory T cell (Treg) cluster (Cluster 0) as the most abundant (approximately 25% of the total Spike-specific CD4+ T cells). This cluster was characterized by the enrichment for gene signatures and genes associated with Treg (e.g., *FOXP3, IKZF2, TIGIT* and *IL2RA*) and by the expression of higher levels of genes encoding MHC class II chains (e.g., *HLA-DR*, -*DP* and -*DQ*). The second most abundant cluster was a Th17-like population (Cluster 1) characterized by increased expression of *CCR6* transcripts and a significant enrichment for Th17 signature genes (**Fig S2A**). These two clusters, along with the clusters representing the classically defined biological subsets, Th1 and follicular helper T cells (Tfh) (Clusters 2 and 3, respectively), accounted for about 78% of all Spike-specific CD4+ T cells. Additional clusters include populations of central memory cells (TCM; Cluster 4), Th2-like cells (Cluster 5) and *FOXP3+* cells that were enriched for signatures linked to follicular regulatory T cells (TFR; Cluster 6) (**Fig S2B**). A less prevalent cluster showed a hybrid gene signature resembling both Clusters 3 and 5 (Tfh/Th2 cells).

**Figure 5.**
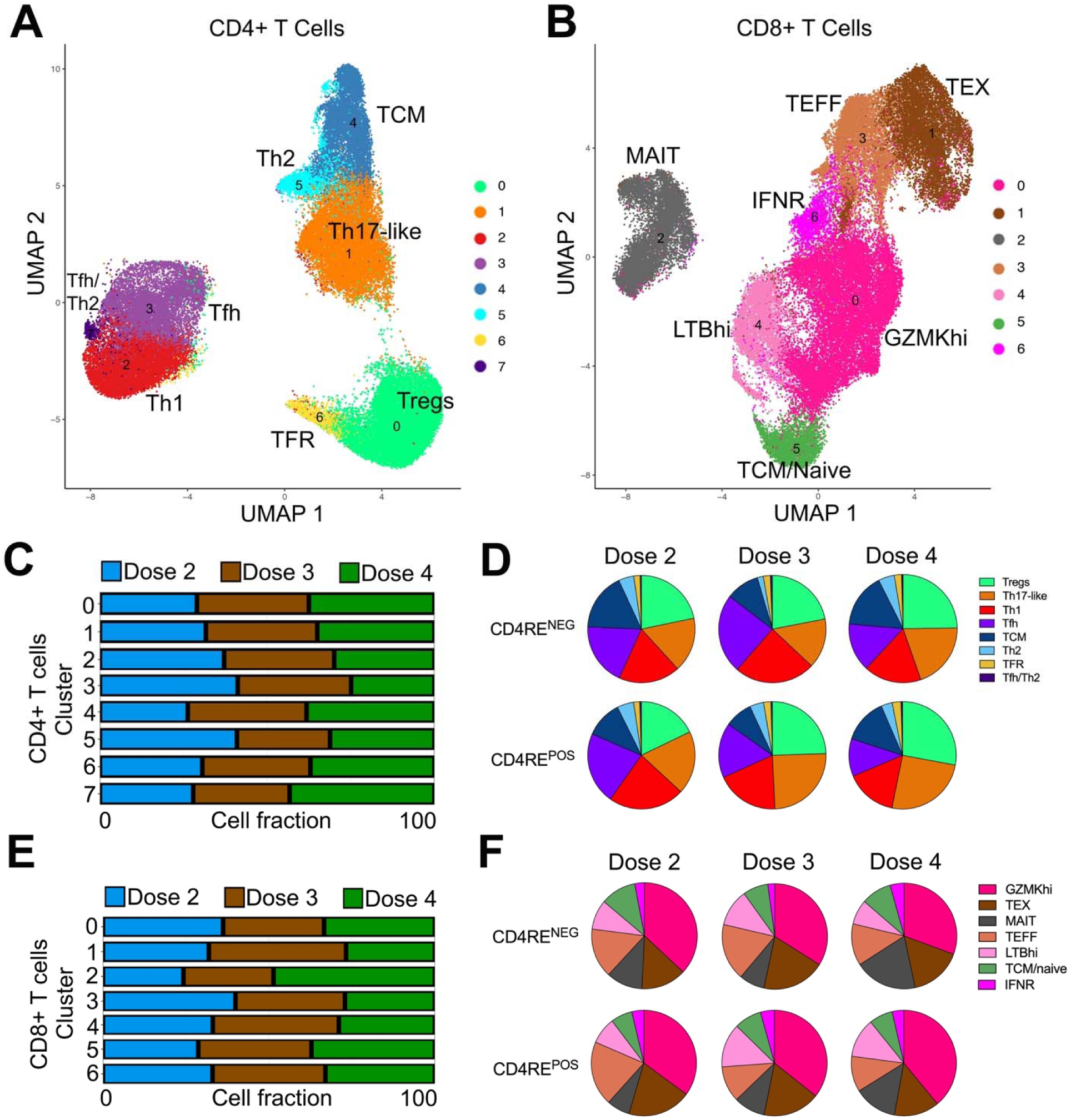
**Spike-Specific CD4+ and CD8+ T Cell subsets are highly diverse and phenotypically stable upon multiple vaccinations**. Single-cell transcriptomes of sorted AIM+ (**A**) CD4+ or (**B**) CD8+ T cells displayed by uniform manifold approximation and projection (UMAP). Cells are colored according to cluster identity as defined by unbiased clustering of 608,720 CD4+ T cells and 460,937 CD8+ T cells. Annotations of cell populations are shown for each cluster. (**C**) Plots show normalized proportion of spike-specific CD4+ T cells in each cluster as function of different vaccine doses. Proportions were normalized according to minimum cell totals across vaccine doses to ensure comparability. (**D**) Pie charts represent the relative frequency of spike-specific CD4+ T cell clusters within each dose cohort in CD4RE^NEG^ (upper panel) or CD4RE^POS^ (lower panel) samples. (**E**) Proportion of spike-specific CD8+ T cells in each cluster as function of different vaccine doses. Proportions were normalized as in (C). (**F**) Pie charts represent the relative frequency of spike-specific CD8+ T cell clusters within each dose cohort in CD4RE^NEG^ (upper panel) or CD4RE^POS^ (lower panel) samples.

**Table 1.** Unbiased cluster annotation of Spike-specific T cell by differential gene expression

| CD4+ T cell Cluster<sup>a</sup> | Description | Markers of interest |
| --- | --- | --- |
| 0 | Tregs | FOXP3, IKZF2, IL1R2, LGALS3, TIGIT, HLA-DRB1, HLA-DPB1, HLA-DPA1, HLA-DRA, HLA-DRB5, ISG20, MX1, IL2RA |
| 1 | Th17-like cells | CCR6, LGALS1, IL4I1, CTSH, ANXA1, ANXA2, VIM, S100A4, S100A6, S100A11, MYL12A, CDKN1A, GZMA, HOPX, LTA, LTB, SELL-negative |
| 2 | Th1 cells | IL2, IFNG, TNF, CSF2, CCL20, IL21, CD200, CD40LG, GZMB |
| 3 | Tfh cells | CXCL13, XCL1, IL3, IL4, IL13, GZMB, ZBED2, DUSP2, DUSP4, PDCD1, CCL20, IFNG |
| 4 | TCM cells | LEF1, KLF2, SELL, TCF7, IL7R, CD27, KLRB1 |
| 5 | Th2 cells | GATA3, IL5, IL13, MX1, ISG15, OAS1, IL17RB, IL10RA, IRF7 |
| 6 | TFR cells | FOXP3, IKZF2, TNFRSF4 (OX-40), TNFRSF9 (4-1BB), TNFRSF18 (GITR), CTLA4, CCR8, TIGIT, ICOS |
| 7 | Tfh/Th2 cells | IL10, CXCL13, IL3, IL4, IL13, GZMB, ZBED2, DUSP2, DUSP4, PDCD1, CCL20, IFNG |
| CD8+ T cell Cluster<sup>b</sup> | Description | Markers of interest |
| 0 | GZMKhi | GZMK, CXCR4, KLF2, KLRC2, KLRC4 (NKG2C and NKG2F), EOMES, GZMM, HLA-DPA1, HLA-DPB1, S1PR4 |
| 1 | TEX cells | GZMB, PRF1, GNLY, IFNG, TNF, CCL4, CCL4L2, CCL3, XCL1, XCL2, CRTAM |
| 2 | MAIT cells | KLRB1, CCL20, IL7R, CXCR6, IFNG, TRAV1-2, TRBV20-1, TRBV6-1 |
| 3 | TEFF cells | GZMB, PRF1, GNLY, IFNG, TNF, NKG7, KLRD1, TNFRSF4, TNFRSF18, LGALS1 |
| 4 | LTBhi cells | LTB, LTA, LGALS1, IFITM1, IFITM2, IFITM3, OAS3, ISG20, IRF1, |
| 5 | TCM/Naive cells | IL7R, TCF7, TCF7, CXCR4, SELL, CD27 |
| 6 | IFNR cells | MX1, MX2, OAS1, OAS3, ISG15, ISG20, IFI6 |

a. Spike-reactive CD4+ T cells
b. Spike-reactive CD8+ T cells

Similarly, a variety of different CD8+ T cell phenotypes were associated with different patterns of gene expression (**Fig 5B** and **Fig S1B**). Out of the seven clusters identified, the one characterized by high expression of granzyme K (*GZMK*)-encoding transcripts was the most abundant (GZMKhi; Cluster 0), encompassing approximately 35% of the total Spike-specific CD8+ T cells. Other prominent clusters, 1 and 3, were enriched for genes linked to effector functions, both showing higher expression of transcripts encoding cytotoxic markers and effector molecules (e.g., *GZMB*, *PRF1*, *GNLY*, *IFNG*, and *TNF*, among others). Notably, Cluster 1 also showed enrichment for gene signatures associated with exhaustion (TEX) (**Table 1** and **Fig S2C**).

An additional cluster exhibited an enrichment for the interferon response (IFNR) gene signature (IFNR cells; Cluster 6), which includes *MX1*, *MX2*, *OAS1*, *OAS3*, *ISG15*, *ISG20*, and *IFI6*. This represents a T cell subset associated primarily with genes related to the immune system’s response to viral infections, constituting a portion of the interferon-stimulated genes (ISGs) activated in response to viral presence (Schneider et al., 2014). A cluster enriched for genes encoding the effector cytokines tumor necrosis factor-α (*LT*α) and lymphotoxin-b (*LT*β) (LTBhi cells; Cluster 4) was also identified, representing a specialized subset sharing similarities with the IFNR subset and potentially contributing to antiviral responses (Schneider et al., 2008; Suresh et al., 2002). We also found a population enriched for naïve/central memory gene signatures (TCM/Naïve cells; Cluster 5) and a population of mucosal-associated invariant T cells (MAIT; Cluster 2), potentially captured by bystander-activation from PBMC stimulation (Kim and Shin, 2019), which were not considered for further analysis.

### Evolution of Spike-specific CD4+ and CD8+ T cell phenotypes as a function of multiple vaccinations

We analyzed the fraction of total response accounted for the various T cell subsets, as a function of the number of vaccinations and CD4RE positivity (**Fig 5C-F**). Across multiple vaccination rounds, the proportions of Spike-specific CD4+ and CD8+ T cell subsets demonstrated remarkable stability (**Fig 5C** and **Fig 5E**, respectively), independent of CD4RE reactivity (i.e., negative vs. positive) (**Fig 5D** and **Fig 5E**).

Most importantly, in longitudinal samples, we did not observe a significant reduction in the expression of multiple effector cytokine transcripts in the most prominent polyfunctional Spike-specific CD4+ T cell subsets (Th1 and Tfh clusters), suggesting that multiple vaccinations do not significantly impair the cytokine expression profile of Spike-reactive CD4^+^ T cells (**Fig 6A**). Similarly, we did not observe a significant reduction in the expression of cytokines or cytotoxicity associated transcripts in effector Spike-specific CD8+ T cells subsets (GZMKhi and TEFF clusters) after repeated vaccinations (**Fig 6B**).

**Figure 6.**
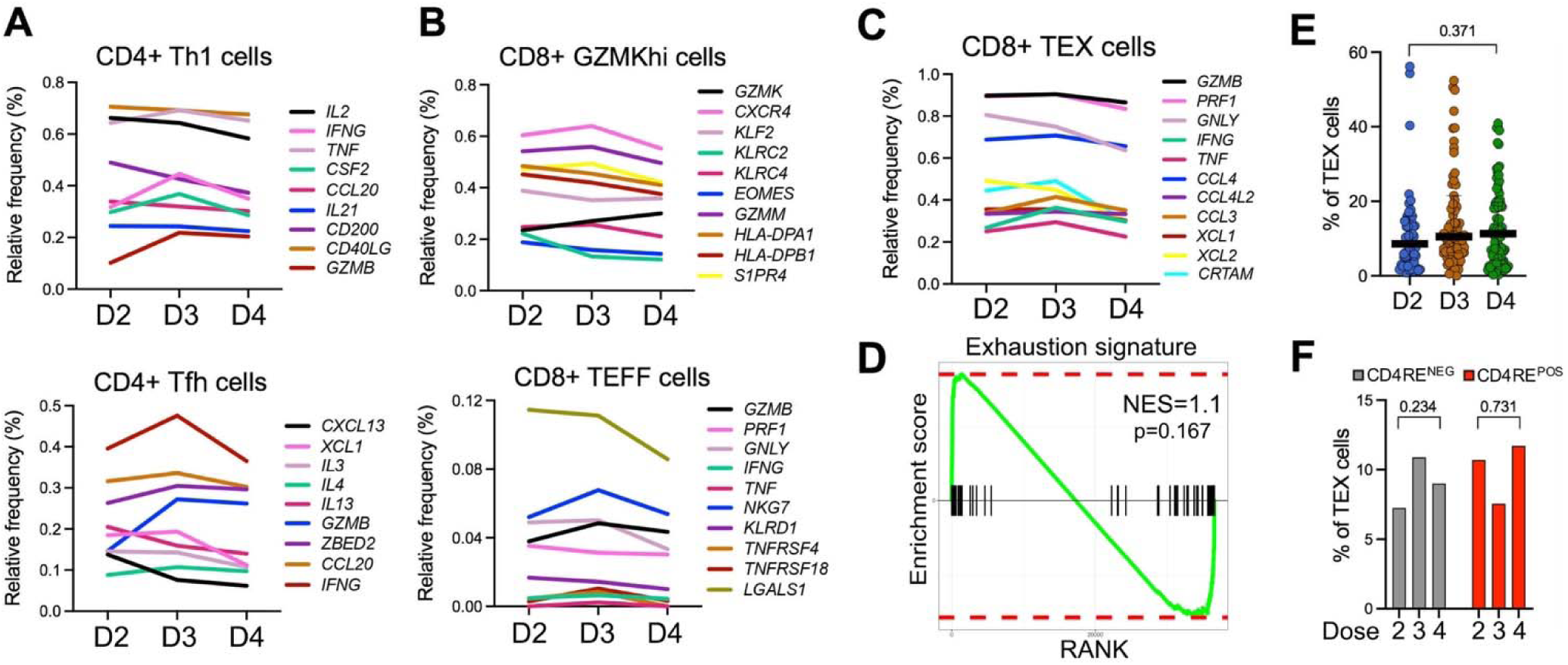
**T cell functionality and exhaustion profiles remain stable after booster vaccination**. Graphs show relative expression of relevant genes cross-sectionally across different vaccine doses for (**A**) CD4+ Th1 cells (cluster 2; top panel) and CD4+ Tfh cells (cluster 3; lower panel), (**B**) CD8+ GZMKhi cells (cluster 0; upper panel), and CD8+ TEFF cells (cluster 3; lower panel) or (**C**) CD8+ TEX cells (cluster 1). Each color-coded gene is indicated. (A-C) Data were analyzed for statistical significance using the Kruskal-Wallis test followed by Dunn’s post-hoc test with FDR correction. (**D**) GSEA for exhaustion signature genes (Exhaustion consensus) across vaccine doses in spike-specific TEX CD8+ T cells. P-value and normalized enrichment score (NES) are indicated. (**E-F**) Graphs represent the relative frequency of CD8+ TEX (cluster 1) cells across timepoints for the indicated vaccine doses in all samples (**E**) or in samples associated with negative or positive CD4RE reactivity (**F**). Data were analyzed for statistical significance using the unpaired Mann-Whitney test. p values are shown.

Analysis of CD8+ T cells gene expression profiles also showed no evidence of T cell exhaustion following multiple COVID-19 vaccinations (**Fig 6C**). Employing Gene Set Enrichment Analysis (GSEA) with a comprehensive exhaustion gene signature derived from nine independent studies of exhausted T cells (e.g. *PDCD1*, *LAG3*, *HAVCR2*, and *TIGIT*, among others) (Kusnadi et al., 2021), we found no significant enrichment for exhaustion markers in the Spike-specific TEX cell population when comparing later timepoints (after D4) to the earlier timepoints following the D2 vaccination (**Fig 6D**). This finding also held true for the proportion of Spike-specific T cells within the TEX cluster (**Fig 6E**). Additionally, the exhaustion signature and proportion of TEX cells remained unchanged regardless of whether individuals had experienced asymptomatic infections (**Fig 6F**).

Despite the overall stability of Spike-specific T cell subset proportions across multiple vaccination rounds, booster vaccination markedly limited the expansion of CD4+ Tfh cells in the peripheral blood one month after vaccination (FC=7.3, 2.5 and 1.9 for D2, D3 and D4, respectively) (**Fig 7A**). The expression of Tfh-associated transcripts encoding *CXCL13* (which promotes germinal center formation) and *CXCR5* (crucial for homing to cells to germinal centers) paralleled the cell frequency dynamics (**Fig 7B,C**). When comparing samples before vaccination and one-month post-vaccination, these transcripts were also notably upregulated (**Fig 7D**). In contrast, expression of *IL2* transcripts, known to suppress the differentiation of Tfh cells, was significantly decreased during this period. We speculate that multiple vaccinations may limit the expansion of circulating Tfh cells in peripheral blood and potentially enhance the maturation and retention of follicular helper T cells in germinal centers or lymphoid organs (Mudd et al., 2022).

**Figure 7.**
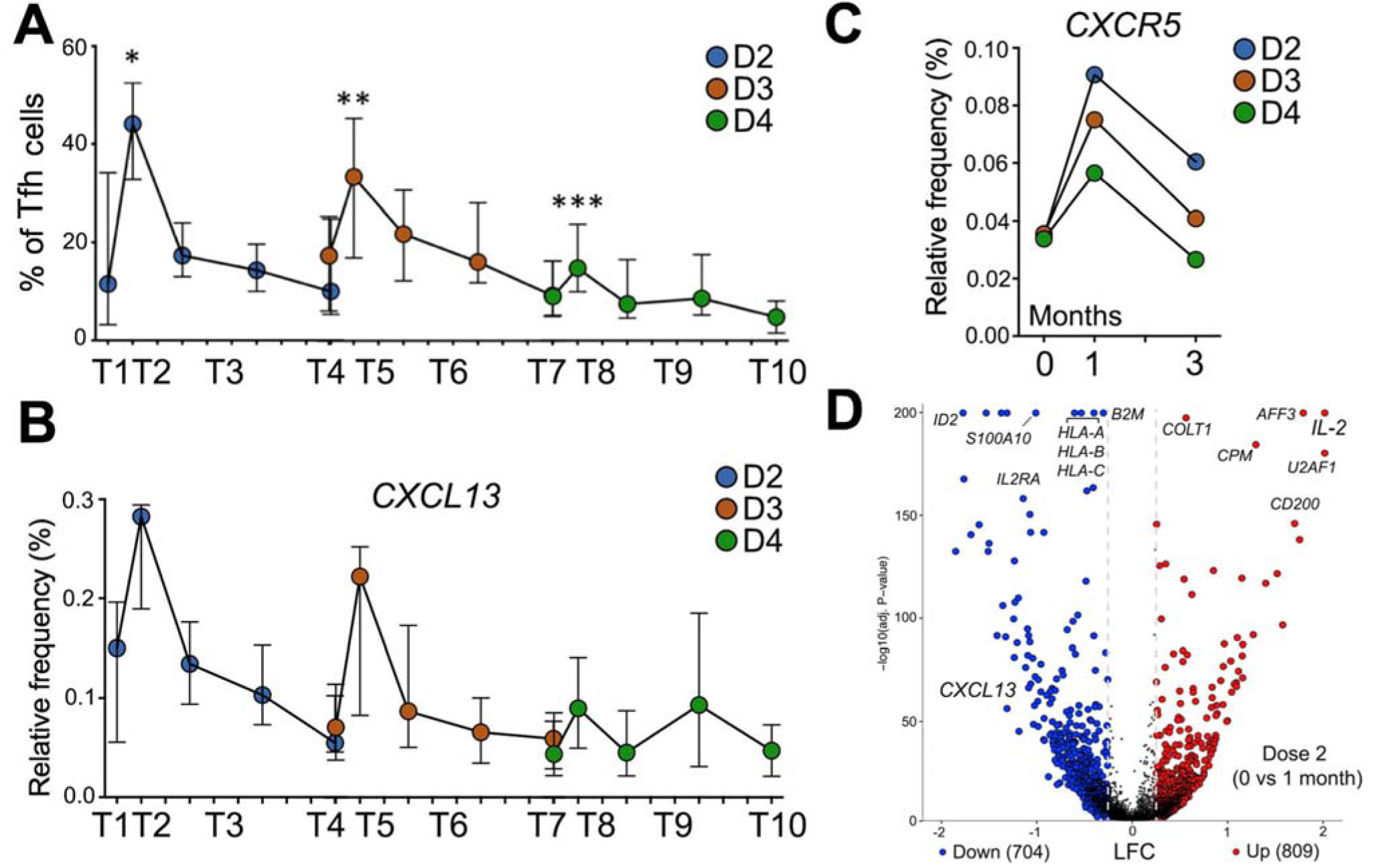
Repeated vaccinations limit peripheral Tfh cell expansion. (**A**) Graphs represent the relative frequency of Tfh cells in multiple vaccinations and timepoints. (**B, C**) Graphs show relative gene expression of (**B**) *CXCL13* and (**C**) *CXCR5* cross-sectionally across different vaccine doses and timepoints (months from vaccination). Color-coded dots represent the median response for each vaccine dose and timepoint. Interquartile range is represented. Data were analyzed for statistical significance comparing timepoints before and after each vaccination, using the unpaired Mann-Whitney test. * p = 0.0056; ** p = 0.0071; *** p = 0.0076. (**D**) Volcano plot shows differentially expressed genes in Tfh cells before (T1) and 30 days after (T2) Dose 2 vaccination. Vertical dotted lines indicate log2 fold change (LFC) cutoff, to call differentially expressed genes. Number of significant genes up- or down-regulated are indicated and dots are color-coded accordingly.

In conclusion, T cell populations and phenotypes were overall stable beyond one month after each booster, with no significant evidence of loss of functionality or progression to exhaustion over multiple vaccinations. Significant changes in frequency were however noted for some clusters, particularly in the context of asymptomatic infections, as detailed in the following paragraphs.

### Asymptomatic infections are associated with increased Th17-like and GZMKhi, IFNR and LTBhi T cell subsets

The data presented above show that previous asymptomatic infection influences the outcome of vaccination in terms of overall magnitude and decay of responses. We similarly reported that previous asymptomatic infection also influences the immunological outcomes of further breakthrough infections (BTIs) (Tarke *et al*., 2024). Accordingly, we explored if CD4RE^POS^ samples would be associated with a qualitative difference in the longitudinal evolution of CD4+ and CD8+ T cell responses. We noted higher frequencies of the Th17-like CD4+ T cell cluster (Cluster 1) in CD4RE^POS^ samples compared to CD4RE^NEG^ samples (**Fig 8A**). This pattern was consistently observed for all different timepoints and vaccine doses (**Fig S3A**). Additional analysis revealed that despite the increase in frequency, there were no significant differences in the gene expression profiles of Th17-like CD4+ T cell cluster from CD4RE^NEG^ and CD4RE^POS^ samples (**Fig S3B**). In the case of CD8+ AIM+ T cells, the IFNR, LTBhi, and GZMKhi subsets were likewise increased in CD4RE^POS^ samples compared to CD4RE^NEG^ samples (**Fig 8B**), at all timepoints and vaccine doses examined (**Fig S3C**). Also, CD8+ GZMKhi, LTBhi, and IFNR T cell subsets from CD4RE^POS^ and CD4RE^NEG^ samples showed minimal differences in their gene expression profiles (**Fig S3D**).

**Figure 8.**
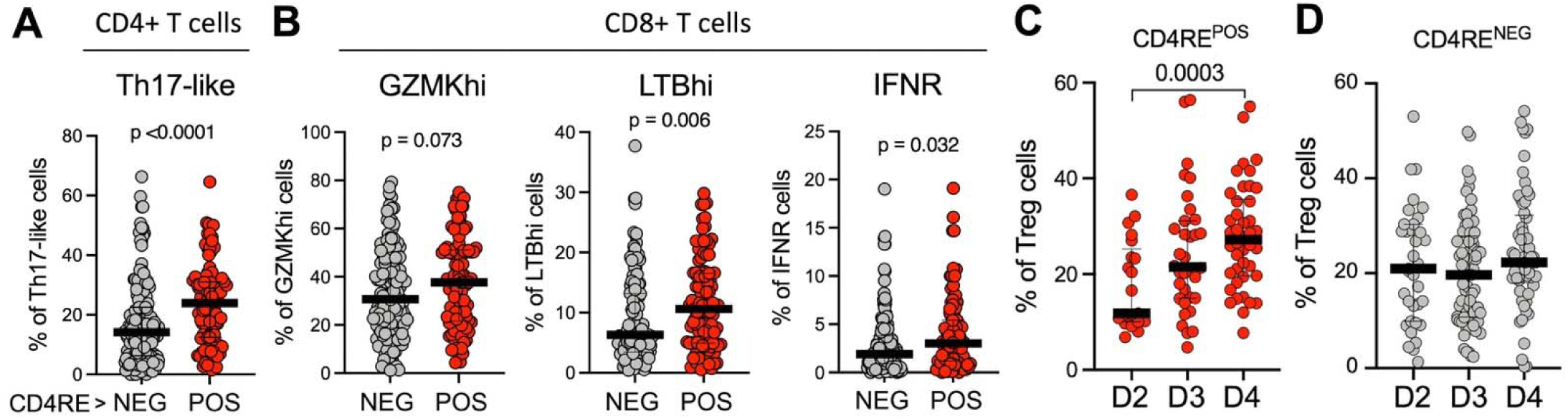
Asymptomatic infections are associated with increased Th17-like and GZMKhi/IFNR cell phenotypes concurrent with Tregs expansion over repeated vaccinations. (**A**) Graphs represent the relative frequency of CD4+ Th17 cells in CD4RE negative and positive samples across doses and timepoints. (**B**) Graphs represent the relative frequency of CD8+ GZMKhi, LTBhi and IFNR cells in CD4RE negative and positive samples across doses and timepoints. (**C, D**) Graph represents the relative frequency of CD4+ Tregs in different vaccine doses across samples associated with (**C**) positive or (**D**) negative CD4RE reactivity. Data were analyzed for statistical significance using the unpaired Mann-Whitney test. p values are shown.

In summary, these results suggest that Spike-specific T cells from samples associated with asymptomatic infections have increased frequency of Th17-like and GZMKhi/LTBhi/IFNR cells, and that the transcriptomic profile of these subsets largely resemble that of the corresponding clusters from vaccinated samples only. These findings are consistent with the observation of higher Th17 and IFN gene signatures in patients with COVID-19 (Daamen et al., 2022; Galbraith et al., 2022; Jovanovic et al., 2023).

### Asymptomatic infections are associated with an increase in Treg cells over repeated vaccinations

As shown above, asymptomatic infections are associated with increased representation of clusters associated with inflammatory and viral effector function.

Notably, the most striking longitudinal change observed with sequential booster vaccinations was a steady increase in the frequency of Spike-specific Tregs, which was only observed when considering CD4RE^POS^ samples (**Fig 8C**) and not when considering CD4RE^NEG^ samples (**Fig 8D**). Interestingly, in the same group of CD4RE^POS^ samples, this kinetic was not observed on TFR cells (Cluster 6), the other discernable sub-population of Tregs (**Fig S4A**). Surprisingly, the increase in Treg frequency over multiple booster vaccinations in CD4RE^POS^ samples was not paralleled by an increase in expression of genes encoding for cytokines or receptors with classic suppressive activity (e.g., *IL10*, *TGFB1*, *CTLA4*, or *TIGIT*) (**Fig S4B**), suggesting that repeated vaccinations do not alter the functional characteristics of Tregs.

In summary, repeated vaccinations in asymptomatic cases are associated with declining Tfh and rising Treg cell populations that could potentially be linked to protection from immunopathology, which goes in line with our previous observation of reduced proportions of SARS-CoV-2-specific Treg subsets in COVID-19 hospitalized patients when compared to non-hospitalized patients (Meckiff et al., 2020).

## DISCUSSION

Our longitudinal analysis of SARS-CoV-2-specific responses following multiple mRNA vaccinations addresses durability, magnitude, and quality of adaptive immune responses and also unveils how unreported, and likely asymptomatic, infections influence the outcome of vaccinations. We find that repeated vaccination increases the magnitude and durability of antibody responses. In contrast, for T cell responses at the 1-3 months timepoints, the magnitude and durability remain relatively unaffected by repeated boosters, as T cell responses quickly plateau and remain stable thereafter. While repeated boosters have little discernible effect on the magnitude of T cell responses, they do affect the quality of T cell responses, particularly in individuals associated with a previously unreported asymptomatic infection.

The incremental enhancement of magnitude and durability of antibody responses with each booster dose observed in this study aligns with recent findings by Srivastava *et al* (Srivastava *et al*., 2024). In our study we further report that asymptomatic infections contributed to more stable antibody responses, potentially reflecting an additional immune stimulation event, and underscoring the complex interplay between vaccination and natural exposure in shaping humoral immunity (Pusnik et al., 2023; Sette and Crotty, 2022). In contrast to antibody kinetics, T cell responses exhibited rapid plateauing followed by remarkable stability, irrespective of antigen exposure through vaccination or asymptomatic infection.

In our study, we detected a substantial prevalence of CD4RE reactivity (27.2%) among vaccinated individuals who had not reported symptomatic infections or positive antigen or PCR SARS-CoV-2 tests at any time during this study. A significant impact of undetected infections on the boostability of immune response outcomes has also been recently reported (Tarke *et al*., 2024). The detection of CD4RE reactivity in approximately one third of samples from participants in this study underscores the limitations of conventional surveillance methods and the likely underestimation of community exposure to SARS-CoV-2. This observation is also consistent with reports that indicate that a significant percentage of SARS-CoV-2 infections, particular of the Omicron lineage are asymptomatic (Tarke *et al*., 2024; Wang *et al*., 2023; Yu *et al*., 2022c) further suggesting that T cell responses may be a more sensitive indicator of prior SARS-CoV-2 exposure than antibody-based assays (Le Bert and Samandari, 2024; Yu *et al*., 2022b). Our results underscore the critical need for proper sample segregation and accurate determination of infection history (Machkovech et al., 2024).

Our scRNA sequencing analysis demonstrates consistent phenotypic stability in both CD8+ and CD4+ T cell subsets across multiple vaccinations, dispelling hypotheses of progressive T cell exhaustion or functional impairment after repeated boosters. This finding is consistent with recent studies by Keeton *et al* (Keeton et al., 2023) and Gray-Gaillard *et al* (Gray-Gaillard et al., 2024), which demonstrated the plasticity and functional resilience of SARS-CoV-2-specific T cells in the context of repeated antigen exposure. Our data also reveal a highly diverse landscape of Spike-specific T cell subsets, notably featuring prominent populations of Tfh cells and regulatory cells. Surprisingly, booster vaccinations appear to limit Tfh cell expansion in peripheral blood, potentially due to their retention in germinal centers (Mudd *et al*., 2022). Conversely, in participants with positive CD4RE reactivity, Treg cells progressively increased as a function of multiple vaccinations. Davis-Porrada *et al* (Davis-Porada et al., 2024) found high frequencies of activated Tregs and regulatory functional profiles that comprised the majority of Spike CD4+T cells in blood and also tissues. Meckiff *et al* (Meckiff *et al*., 2020) reported that SARS-CoV-2-specific Tregs are reduced in hospitalized COVID-19 patients, and Franco *et al* (Franco et al., 2023) reported that vaccine-induced Tregs may play a crucial role in preventing excessive inflammation. Our observation of the emergence of a regulatory population following repeated vaccinations in individuals with asymptomatic infection suggests that mRNA vaccines may elicit a balanced immune response, potentially mitigating the risk of SARS-CoV-2 immune-mediated pathology.

Interestingly, despite the overall increase in Tregs, Tfh retention or even slight increases of effector T cell function, we observed a significant increase in the proportion of Th17-like cells in individuals with evidence of asymptomatic infection. These Th17- like cells could have a more beneficial role in the control of antiviral activity in the context of asymptomatic infections. Our analysis of CD8+ T cell subsets also revealed that the populations of GZMKhi and IFNR cells were increased in individuals with potential asymptomatic infections. Since Spike-specific CD8+ T cell response is known to correlate with viral clearance and to predict COVID-19 progression (Kedzierska and Thomas, 2022; Koutsakos et al., 2023; Zhang et al., 2023), it is possible that these particular subsets could be associated with increased viral control and better outcomes in COVID-19 patients.

In conclusion, our study provides a comprehensive picture of the evolving adaptive immune landscape following repeated SARS-CoV-2 mRNA vaccinations. Booster vaccinations lead to stable memory CD4+ and CD8+ T cell populations without inducing cell exhaustion, and our data further suggests that asymptomatic SARS-CoV-2 infection may be associated with expansion of Tregs, potentially providing a balance between effector T cell function facilitating viral immunity and regulatory functions limiting immunopathology.

### Limitations of the Study

Our study is limited to analysis of peripheral blood responses. Future studies will address the features of tissue-resident T cells and in particular mucosal immune responses, following methodologies recently described (Davis-Porada *et al*., 2024; Ramirez et al., 2024). By design, our study included only individuals that did not report symptomatic infections. Future studies will specifically address the characteristics of vaccinated individuals that experienced symptomatic breakthrough infections, specifically at the level of scRNA profiles and T cell subsets. Due to our study design excluding vaccinated individuals with symptomatic breakthrough infections, we did not perform scRNA analysis of T cell responses to non-Spike antigens, which might differ from Spike-specific T cell responses. Extending this study with integrated analyses of TCRseq data and in BTI cohorts can provide valuable insights into the effects of multiple booster vaccinations and infections on the dominance of T cell clonotypes, further enhancing our understanding of immune protection against SARS-CoV-2.

## RESOURCE AVAILABILITY

### Lead Contact

Further information and requests for resources and reagents should be directed to the lead contact, Dr. Alessandro Sette.

### Materials Availability

Peptide pools used in this study will be made available to the scientific community upon request, and following execution of a material transfer agreement, by contacting Dr. Alessandro Sette.

### Data and Code Availability

The published article includes all data generated or analyzed during this study, and summarized in the accompanying tables, figures and supplemental materials. Code written for data analysis and visualization is available in our repository on GitHub (https://github.com/vijaybioinfo/BOOSTER_2025). Sequencing data for this study is being deposited to the Gene Expression Omnibus and an accession number will be provided upon completion. Any additional information required to reanalyze the data reported in this work paper is available from the Lead Contact upon request.

## ACKNOWLEDGEMENTS

We wish to acknowledge all subjects for their participation and for donating their blood and time for this study. We are grateful to Gina Levi and the La Jolla Institute for Immunology clinical core relentless efforts in sample coordination and blood processing. We thank the Flow cytometry facility at La Jolla Institute for Immunology for technical assistance with cell sorting. This project has been funded in whole or in part with Federal funds from the National Institute of Allergies and Infectious Diseases, National Institutes of Health, Department of Health and Human Services, under Contract No. 75N93019C00065, Grants No. U19 AI142742 and U19 AI118626.

## AUTHOR CONTRIBUTIONS

Conceptualization, R.d.S.A., and A.S.; methodology, V.F.R., E.D.Y., R.I.G., A.A., E.A.E., A.M.P., E.J., and B.G.; formal analysis, R.d.S.A., V.F.R., R.I.G., A.A., A.M.P and B.G.; investigation, R.d.S.A., V.F.R., P.V., and A.S.; support of investigation, A.F., J.M.D., S.C., G.S., and D.W.; funding acquisition, P.V., and A. S.; writing, R.d.S.A., V.F.R., P.V., and A.S.; supervision, R.d.S.A., J.M.D., G.S., D.W., P.V., and A.S.

## MATERIAL AND METHODS

### EXPERIMENTAL MODEL AND SUBJECT DETAILS

We conducted a 3-year observational study (January 2021 to January 2024) following several hundred participants. Our analysis focused on 78 vaccinated participants, healthy adult donors who did not report COVID-19-related symptoms at any timepoint. Participants were rigorously screened to ensure they tested negative for SARS-CoV-2 both before and during the sample collection period. The Clinical Core at the La Jolla Institute collected all samples. All donors were from San Diego, CA and provided informed consent with sample collection approved by the LJI Institutional Review Board under IRB Protocol # VD-214. Adults of all races, ethnicities, ages, and genders were eligible to participate. Each participant was assigned a study identification number with clinical information recorded. In order to ensure a uniform group of donors, individuals who received bivalent booster shots or adhered to a different time regimen for the first two doses, classified as full vaccination, were excluded from the study. The study participants’ median age was 44 years (21-81 range), with an overall 67% female and 33% male overall composition. The predominant race was Caucasian accounting for 67% of the total cohort. Moderna (mRNA-1273) and Pfizer (BNT162b2) vaccines (administered in 77/78 participants) were fairly evenly distributed, with 50.0% and 48.7% for the full vaccination (after 2 shots), 56.5% and 42.0 % for the first booster (after 3 shots), and 44.8% and 53.5% for the second booster (4 shots), respectively. A single donor that received 4 different doses of Novavax (NVX-CoV2373) accounted for the rest. Demographics reflect the overall characteristics of the volunteers who are enrolled in LJI SARS-CoV-2 studies (**Table S1**).

### Sample collection times and sub-group organization

Samples associated with each visit were organized according to the temporal window in which they were collected: within than 30 days after full vaccination (T2), within than 30 days post-first booster (T5), and within than 30 days post-second booster (T8). The longitudinal intervals between vaccinations were further segmented into the periods of 1-6 months post-full vaccination (T3, T6, and T9) or 6-12 months post-full vaccination (T4, T7, and T10). The median time interval preceding full vaccination was 30.5 days for the pre-vaccination period, while post-vaccination samples were collected at a median interval of 18.5 days after full vaccination (T2). Regarding the first boost vaccination (T5), the median pre-vaccination interval was 7.5 days, and post-vaccination samples were obtained at a median interval of 16 days. For the second booster vaccination, samples were collected with a median of 11.5 days before vaccination and 25.5 days post-vaccination.

## METHOD DETAILS

### Peripheral blood mononuclear cells (PBMCs) and plasma isolation

The blood samples were collected in heparin-coated blood bags. Peripheral blood mononuclear cells (PBMCs) were isolated using the same methodology as in previous studies (Dan et al., 2021b; Grifoni et al., 2020a; Zhang et al., 2022b), employing Ficoll-Paque (Lymphoprep, Nycomed Pharma, Oslo, Norway). In brief, whole blood was centrifuged for 15 minutes at 400g to separate the cellular fraction and plasma. Plasma was collected and stored at -20°C. PBMCs were separated by density-gradient sedimentation with Ficoll-Paque® PLUS. Isolated PBMCs were washed twice with RPMI 1640 to remove any residual Ficoll-Paque® PLUS. Subsequently, the PBMCs were resuspended in freezing media containing 90% fetal bovine serum (FBS; Hyclone Laboratories, Logan, UT) and 10% dimethyl sulfoxide (DMSO; Gibco, Waltham, MA) before time and temperature-controlled cryopreservation in liquid nitrogen until used for experiments.

### SARS-CoV-2 ELISAs

The SARS-CoV-2 ELISAs were performed as described in previous studies (Dan *et al*., 2021b; Zhang *et al*., 2022b). Briefly, Corning 96-well half-area plates (ThermoFisher 3690) were coated with 1 μg/ml of SARS-CoV-2 RBD Spike protein, followed by overnight incubation at 4°C. Subsequently, plates were blocked with 3% milk in phosphate-buffered saline (PBS) containing 0.05% Tween-20 for 1.5 hours at room temperature. Plasma was heat-inactivated at 56 °C for 30 to 60 minutes, then diluted in 1% milk containing 0.05% Tween-20 in PBS, starting at a 1:3 dilution followed by serial dilutions of three, and incubated for 1.5 hours at room temperature. Plates were washed five times with 0.05% PBS-Tween-20. The secondary antibody, an anti-human IgG peroxidase antibody produced in goat (Sigma A6029), was used at a 1:5,000 dilution and diluted in 1% milk containing 0.05% Tween-20 in PBS. Plates were read on a Spectramax Plate Reader at 450 nm using the software SoftMax Pro. The limit of detection (LOD) was defined as 1:3. The limit of sensitivity (LOS) was established based on uninfected subjects using pre-pandemic plasma.

### SARS-CoV-2 Megapools (MP)

To study T cell responses against SARS-CoV-2, two peptide pools (Megapools; MP) were prepared following the MP approach, previously outlined as a comprehensive method for analyzing T cell responses across diverse epitopes and populations (da Silva Antunes *et al*., 2023). A MP of 15-mer peptides overlapping by 10 spanning the entire Spike protein sequence (253 peptides) and a MP (CD4RE) composed of 284 experimental defined epitopes from non-Spike (R) region of SARS-CoV-2 were selected as previously described (Grifoni *et al*., 2020a; Yu *et al*., 2022b). The CD4RE MP was further employed to identify previous SARS-CoV-2 infection (Yu *et al*., 2022b). All peptides were synthesized as crude material (TC Peptide Lab, San Diego, CA), individually resuspended in DMSO at a concentration of 20 mg/mL. MPs were prepared by pooling individual peptides followed by lyophilization (da Silva Antunes *et al*., 2023), and resuspended at 1 mg/mL in DMSO.

### Activation Induced Marker (AIM) assay and flow cytometry analysis

To detect T cell-specific responses we employed an AIM assay methodology in combination with peptide pool stimulation using the dual activation marker expression of OX40+4-1BB+ or CD69+4-1BB+ to detected antigen-specific CD4+ and CD8+ T cells, respectively (da Silva Antunes *et al*., 2023; Dan *et al*., 2016). Briefly, PBMC vials were thawed with subsequent determination of cell number. PBMCs were then plated in 96-well U-bottom plates at a concentration of 1-2×10^6 cells per well and immediately cultured together with MPs (1 µg/mL), or phytohemagglutinin-L (PHA) (10 µg /mL; Roche, San Diego, CA) and DMSO as positive and negative controls, respectively, in 5% human serum (Gemini Bio-Products) for 18-24 h. After culture, cells were collected, washed and antibody staining performed as previously described (Zhang *et al*., 2022a). A comprehensive list of antibodies utilized is shown in **Table S4**. All samples were acquired on a Cytek Aurora (Cytek Biosciences, Fremont, CA) and analyzed with FlowJo 10.9 software (Tree Star, Ashland, OR). The gating strategy is illustrated in **Fig S5**.

The data were normalized with a minimum response level set at 0.005%. The specific T cell responses were calculated by subtracting the background (DMSO stimulation) values. For each population, the limit of detection (LOD) was determined as the upper 95% confidence interval of the DMSO values, while the limit of sensitivity (LOS) was calculated as the median plus 2 times the Standard Deviation (SD) of DMSO. The Stimulation Index (SI) was calculated as the percentage of stimuli response divided by the percentage of response in the DMSO control. Positive responses were defined as responses greater than LOS, with SI greater than 2 for CD4+ T cells or SI greater than 3 for CD8+ T cells. Responses with SI lower than 2 for CD4+ T cells or SI lower than 3 for CD8+ T cells were normalized to LOD. Donors associated with undetected BTIs were identified by the first timepoint at which their CD4RE MP reactivity responses surpassed a highly stringent 10-fold Stimulation Index threshold.

### AIM+ T cell sorting and sample preparation

Following the previously described protocols, PBMCs were thawed, stimulated with Spike antigen peptide pools for 24 hours, washed and stained with an antibody cocktail containing CD8 BUV496 (1:50; BioLegend, clone RPA-T8, Cat# 612942), CD3 BUV805 (1:50; BioLegend, clone UCHT1, Cat# 612895), CD14 V500 (1:50; BioLegend, clone M5E2, Cat# 561391), CD19 V500 (1:50; BioLegend, clone HIB19, Cat# 561121), CD4 BV605 (1:100; BioLegend, clone RPA-T4, Cat# 562658), CD69 PE (1:50; BioLegend, clone FN50, Cat# 555531), OX40 PE-Cy7 (1:50; BioLegend, clone Ber- ACT35, Cat# 350012), 4-1BB APC (1:25; BioLegend, clone 4B4-1, Cat# 309810), Live/Dead Viability dye eFLuor506 (0.1:100; Invitrogen, Cat# 65-0866-14), brilliant staining buffer and PBS before being refrigerated for 30 minutes at 4°C, protected from light. TotalSeqTM-C oligonucleotide-conjugated antibodies (BioLegend) were also added at this step at 0.01mg/mL final concentration (one distinct antibody per sample). After two washes in PBS, cells were resuspended into 500 μl of MACS buffer (PBS containing 2mM EDTA (pH 8.0) and 0.5% BSA) and stored at 4°C until flow cytometry acquisition. Activated CD4+ (OX40+4-1BB+) and CD8+ (4-1BB+CD69+) AIM+ cells were subsequently sorted using a BD FACSAria Fusion cell sorter (Becton Dickinson), and by employing the gating strategy represented in **Fig S5**. After sorting, ice-cold PBS was added, cells spun down, and single-cell libraries prepared as described below.

### Cell isolation and single-cell RNA-seq library preparation

For single-cell RNA-seq assays (10x Genomics, Pleasantown), in average 20,000- 50,000 AIM+ T cells per subject were sorted directly into low-retention, sterile 1.5 ml collection tubes (Thermo-fisher) pre-chilled on ice and containing 500 µl of PBS:FBS solution (1:1, vol:vol) supplemented with RNAse inhibitor (1:100, Takara Bio). 5 to 6 different samples were multiplexed using the TotalSeq-C DNA-oligo barcoded cell-surface housekeeping molecule antibodies (Biolegend) totaling approximately 60,000 cells per 10X lane (equivalent to one well of the 10X chip). Multiplexed samples were centrifuged at 600 ×g for 10 minutes at 4°C, and the supernatant was carefully removed, leaving behind 10-12 µl of residual volume. Cell pellets were resuspended in 25 µl of 10x Genomics-compatible resuspension buffer (0.22 µm filtered PBS containing 0.04 % ultrapure bovine serum albumin, Sigma-Aldrich). A 33 µl aliquot of the resuspended cells was transferred to an 8-strip PCR tube for downstream processing as per the 10x Genomics protocol. Left-over was used as cell-counting quality control check.

Following the manufacturer’s recommendations, single-cell RNA libraries were generated using the 10x Genomics standard 5’TAG v2 chemistry. Both cDNA amplification and library preparation were carried out using 13 PCR cycles. Barcoded cDNA products were collected, quantified, and pooled at equimolar concentrations. The libraries were sequenced on the Illumina Novaseq6000 (NIH, S10) platform with paired-end sequencing (S4 100x100cycle, Illumina) configured as follows: read length: read 1, 100 cycles; read 2, 100 cycles; i7 index, 10 cycles and i5 index 10 cycles.

Size and quantity quality controls were performed throughout the procedure by capillary DNA high sensitivity electrophoresis (HS NGS Fragment Kit,1-6000bp, Fragment Analyzer, Agilent) and Picogreen assay (Quant-iT™ PicoGreen™ dsDNA Assay Kits and dsDNA Reagents). Sequencing depth was aimed to reach 30,000 reads per cell for gene expressions, 8,000 reads/ cell for TCR, and antibody-feature sequencing (multiplexing analysis). In total for this project, we sequenced around 683,250,903 reads with 42,000 mean read per cell.

### Single-cell transcriptome analysis

Data from scRNA-seq libraries were preprocessed by employing the 10x’s cellranger suite (v7.1.0) and the pre-built GRCh38 human genome reference (2020-A) as previously described (Kusnadi *et al*., 2021; Meckiff *et al*., 2020). Gene expression data from individual libraries were merged into one aggregated set for CD4+ T cells and another for CD8+ T cells. Metric reports for all single-cell transcriptome libraries and aggregated datasets, provided in **Table S5**, indicate good-quality data.

As detailed before (Kusnadi *et al*., 2021; Meckiff *et al*., 2020), the Seurat toolkit (v3.2.3) (Stuart et al., 2019) was employed in R to perform quality control (QC), unbiased clustering, dimensionality reduction and cluster annotation on each aggregated dataset. The following QC criteria were enforced to minimize doublets and eliminate low-quality transcriptomes: 1,500 ≥ unique molecular identifier (UMI) count ≤ 20,000; 800 ≥ gene count ≤ 4,400; and mitochondrial UMI percentage ≤ 10%. Hashtag oligonucleotide (HTO) data were leveraged to perform sample deconvolution and further identify and remove multiplets, with the HTO data processed with the cellranger suite (v7.1.0) and deconvolution performed with the function *MULTIseqDemux* function from Seurat (v3.2.3) as detailed before (Kusnadi *et al*., 2021; Meckiff *et al*., 2020). For dimensionality reduction and unbiased clustering, only the most variable genes accounting for 30% of the total standardized variance and the 30 principal components (PCs) were considered as arguments in the workflow described previously (Kusnadi *et al*., 2021; Meckiff *et al*., 2020). The final set of clusters identified at resolutions 0.4 and 0.2, respectively, for CD4+ and CD8+ T cells, were considered for in-depth annotation on the basis of their markers and gene set enrichment analysis (GSEA), as well as for downstream analyses. GSEA was performed with the *fgsea* package (Korotkevich et al., 2021) as detailed before (Kusnadi *et al*., 2021; Meckiff *et al*., 2020) with the consensus exhaustion, TH17 and TFR gene signatures generated, respectively, in (Kusnadi *et al*., 2021), (Arlehamn et al., 2014) and (Eschweiler et al., 2021). For comparisons involving early and late time points (**Fig 6D**), samples collected at 1, 3, 6 and 9 months following the second vaccine dose were altogether considered as early time points, whereas samples collected at 3, 6, 9 and 12 months following the fourth vaccine dose were grouped and designated as late time points.

### Single-cell differential gene expression analysis

Transcripts enriched in each T cell population at the indicated resolutions were identified by the function *FindAllMarkers* from Seurat (v3.2.3) (Stuart *et al*., 2019) with MAST employed as the statistical framework for differential expression analysis (Finak et al., 2015). Markers for the T cell subsets described herein are listed in **Table S6**. To compare the transcriptomic profiles of specific T cell subsets between different timepoints, differential gene expression analysis (DGEA) was performed in R with the MAST package (v1.12.0) (Finak *et al*., 2015). A gene was considered as differentially expressed if Benjamini-Hochberg adj. P value < 0.05 and either log_2_ fold change (LFC) ≥ 0.25 or LFC ≤ -0.25.

## QUANTIFICATION AND STATISTICAL ANALYSIS

Statistical analyses were performed in GraphPad Prism 10.0.2, unless otherwise stated. Data plotted in linear scale were expressed as Mean ± Standard Deviation (SD), while the data plotted in logarithmic scales were expressed as Geometric Mean ± Geometric Standard Deviation (SD). Unpaired comparisons between groups were performed using the nonparametric two-tailed, and unpaired Mann-Whitney test or Kruskal-Wallis test adjusted with Dunn’s test for multiple comparisons and FDR correction. Longitudinal comparisons between two paired groups were performed using the nonparametric two-tailed paired Wilcoxon’s test. The estimated t ½ was calculated based on linear mixed effects model using R package nlme as previously described by (Cohen et al., 2021). Spearman’s rank correlation coefficient test was used for association analysis. Details of the statistic method employed and the values pertaining to significance and correlation coefficient (R) are noted in the respective figure, and P<0.05 defined as statistically significant.

---

**Figure S1.**
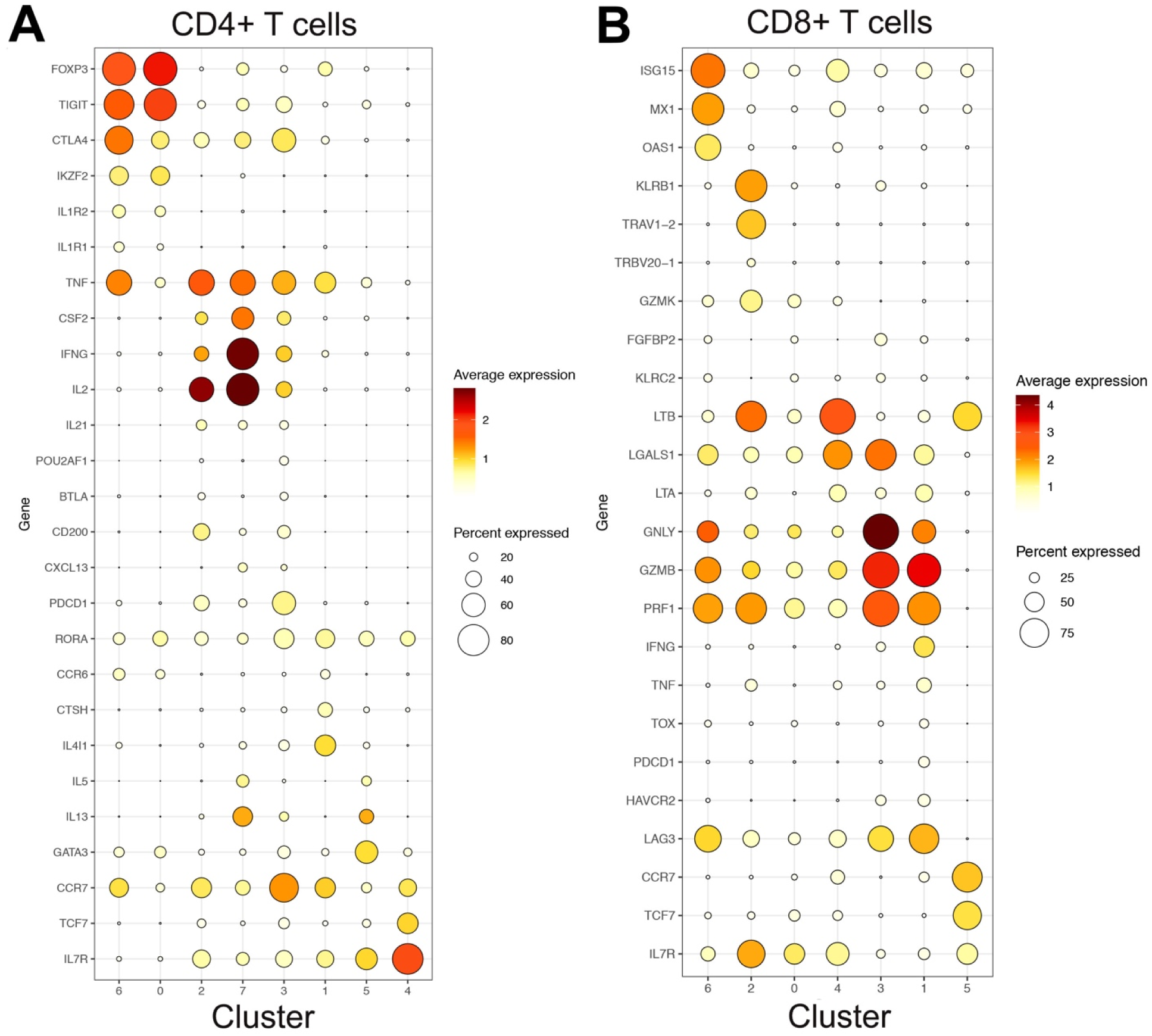
Characterization of Spike-Specific T Cell Subsets. (**A-B**) Dot plots show mean Seurat- normalized expression (color scale) and fraction of marker-positive cells (size scale) for selected markers in each CD4+ T cell cluster (**A**) and in each CD8+ T cell cluster (**B**).

**Figure S2.**
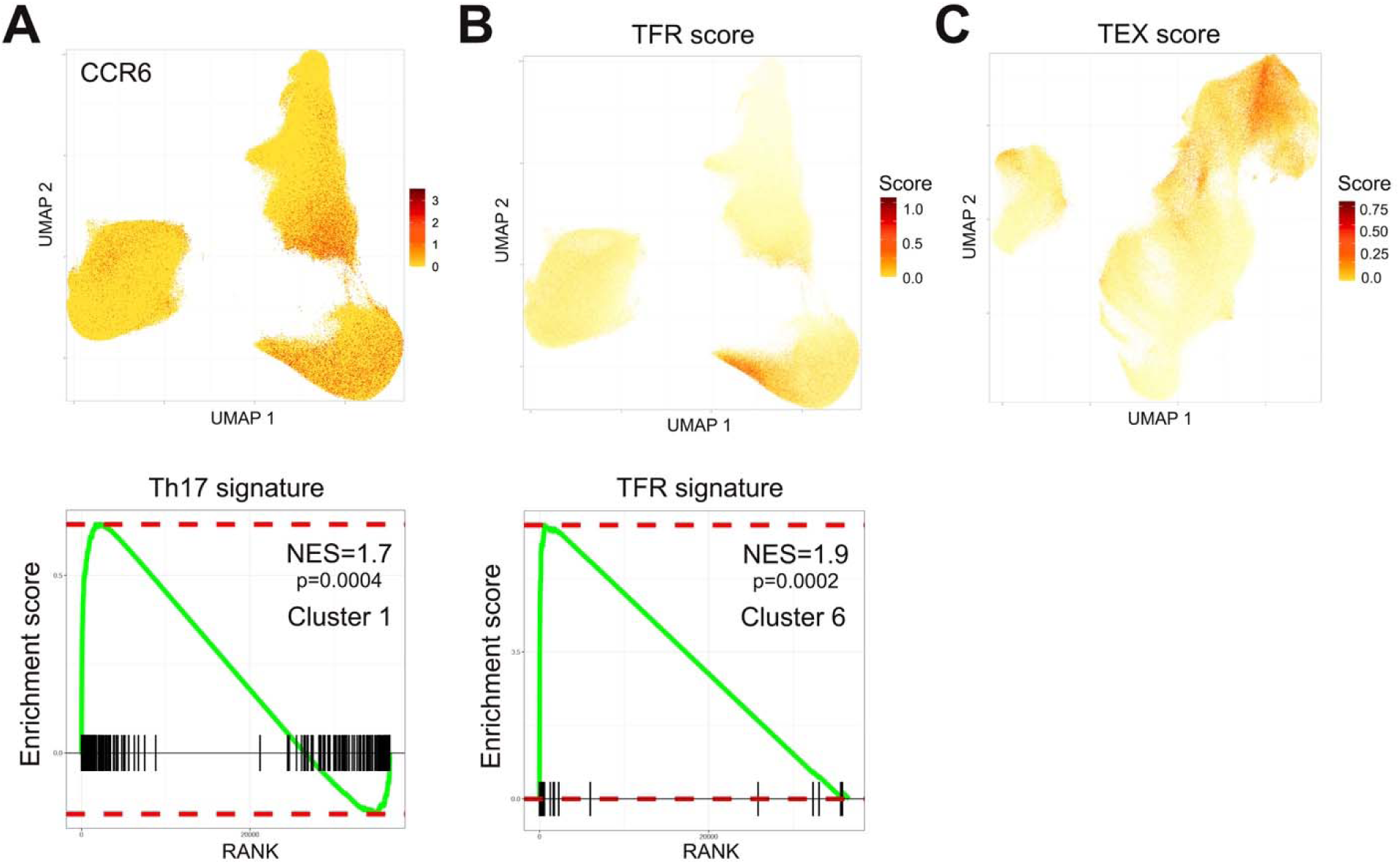
Cluster characterization and definition by gene signature enrichment. (**A**) UMAP showing Seurat-normalized expression of *CCR6* in Spike-specific CD4+ T cells (top), and gene set enrichment analysis (GSEA) plot shows significant enrichment for the Th17 signature in cluster 1 of CD4+ T cells (bottom). (**B**) UMAP showing module scores of the consensus TFR signature in Spike-specific CD4+ T cells (top), and GSEA plot shows significant enrichment for the TFR signature in cluster 6 of CD4+ T cells (bottom). (**C**) UMAP showing module scores of the consensus exhaustion signature in Spike-specific CD8+ T cells. P-value and normalized enrichment score (NES) are indicated in each GSEA graph.

**Figure S3.**
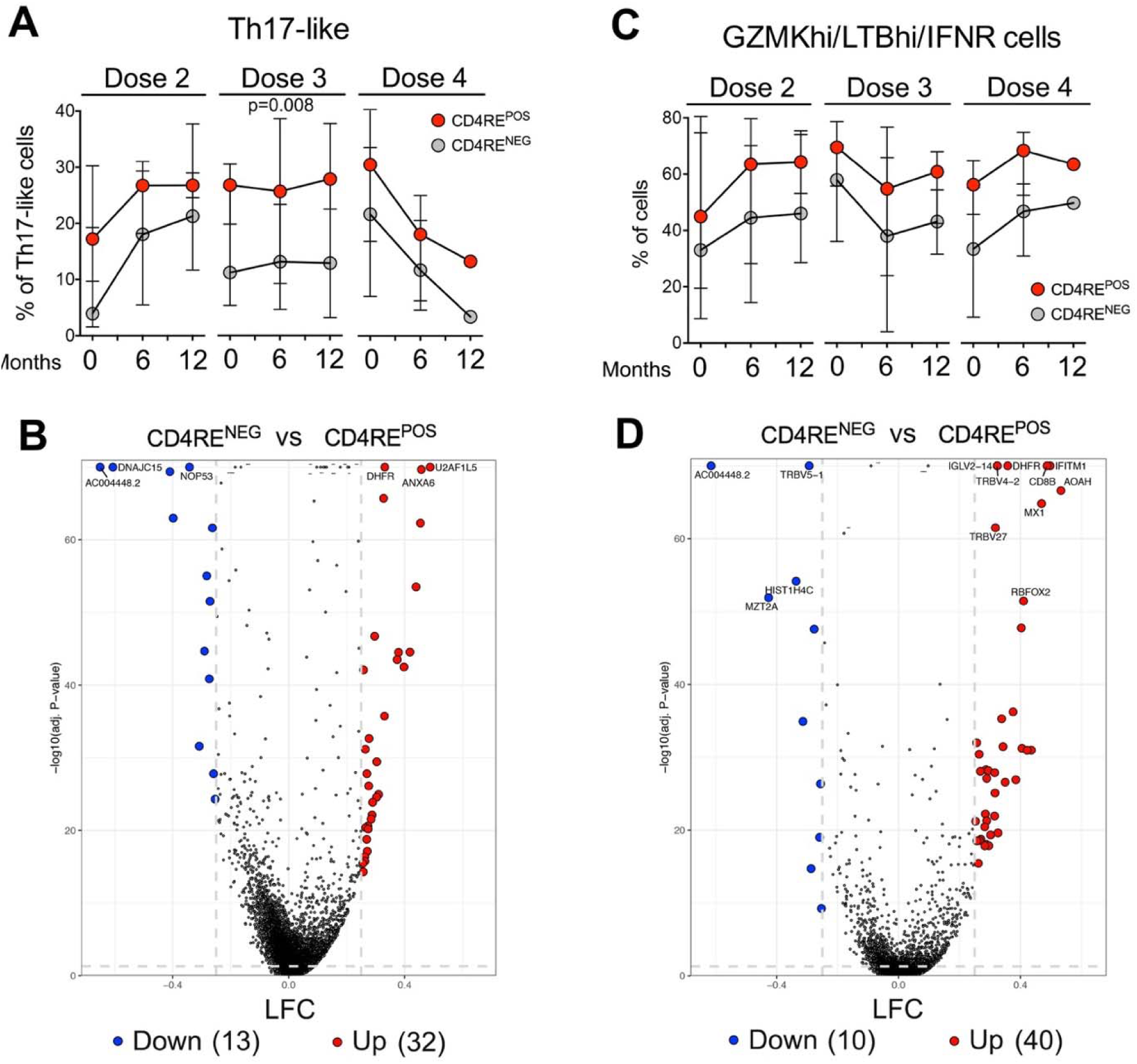
**Kinetics of Th17-like and GZMKhi/IFNR cell phenotypes and gene profile between CD4RE^NEG^and CD4RE^POS^ samples over repeated vaccinations**. (**A**) Graph represent the relative frequency of CD4+ Th17-like cells in CD4RE-negative (grey) and -positive (red) samples and shown for different doses and timepoints (months from each vaccination). Interquartile range is represented. Mann- Whitney test was performed between groups. p value is shown. (**B**) Volcano plot shows genes differentially expressed in CD4+ Th17-like cells between CD4RE-negative and -positive samples across all doses and timepoints. Vertical and horizontal dotted lines indicate cutoffs set, respectively, for p-value and log2 fold change (LFC), to call differentially expressed genes and dots are color-coded accordingly. (**C**) Graphs represent the relative frequency of CD8+ GZMKhi/LTBhi/IFNR clusters (combined) in CD4RE- negative (grey) and -positive (red) samples for different doses and timepoints (months from each vaccination). Interquartile range is represented. (**D**) Volcano plot shows genes differentially expressed in CD8+ GZMKhi/LTBhi/IFNR clusters (combined) between CD4RE-negative and -positive samples across all doses and timepoints. Dots are color-coded as in (B).

**Figure S4.**
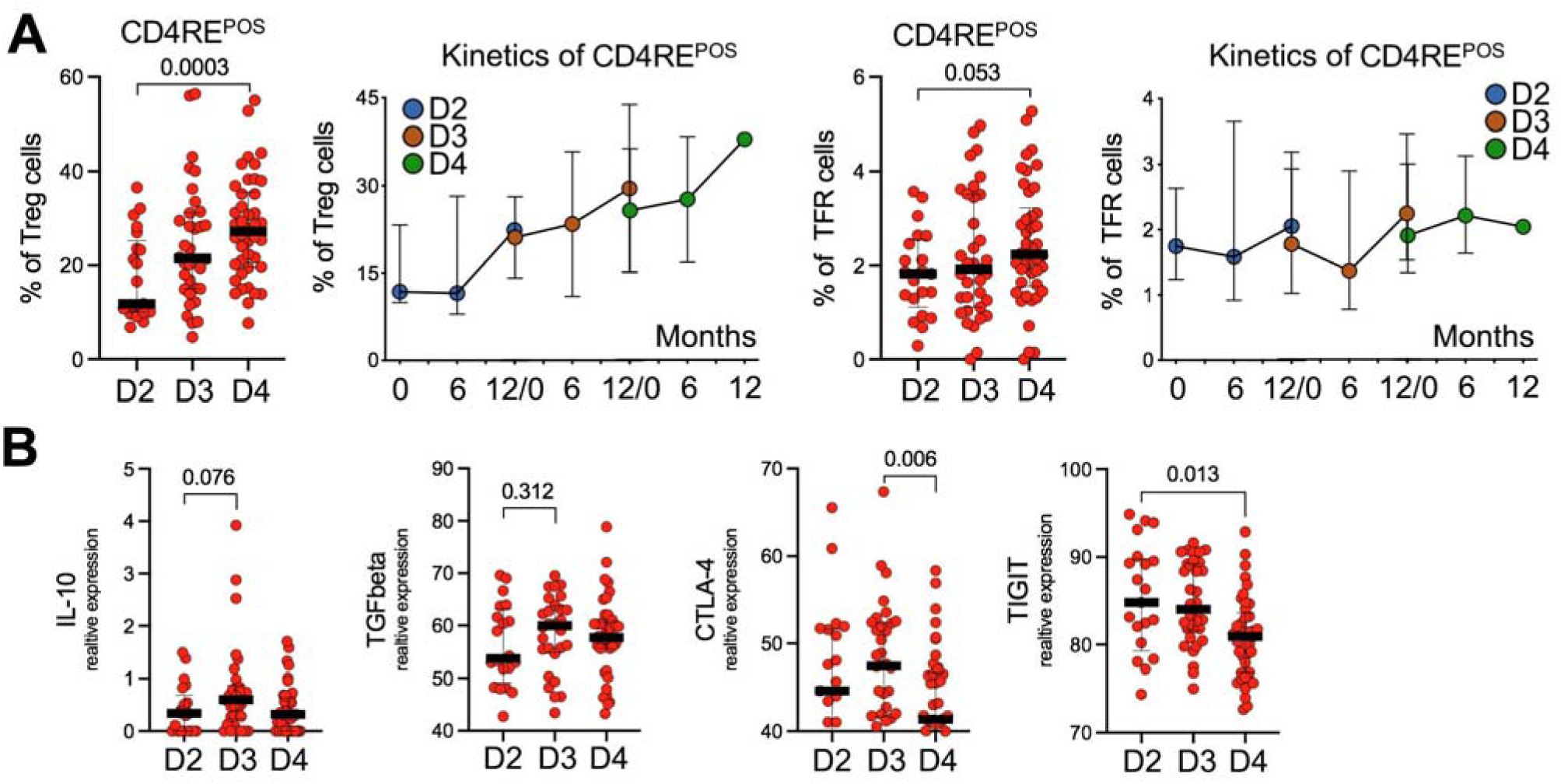
**Kinetics of Tregs and TFR cells and expression of Treg-related suppression genes over multiple vaccinations**. (**A**) Graphs represents the relative frequency of CD4+ Tregs (left) or CD4+ TFR cells (right) in different vaccine doses and timepoints (months from each vaccination) specifically in samples with CD4RE positive responses. Color-coded dots represent the median response for each vaccine dose and timepoint. Interquartile range is represented. (**B**) Graphs show relative frequency of cells positive for selected transcripts (*IL10*, *TGFB1*, *CTLA4* and *TIGIT*) in different vaccine doses. Each red dot represents one individual sample. Data were analyzed for statistical significance using the unpaired Mann-Whitney test. p values are shown.

**Figure S5.**
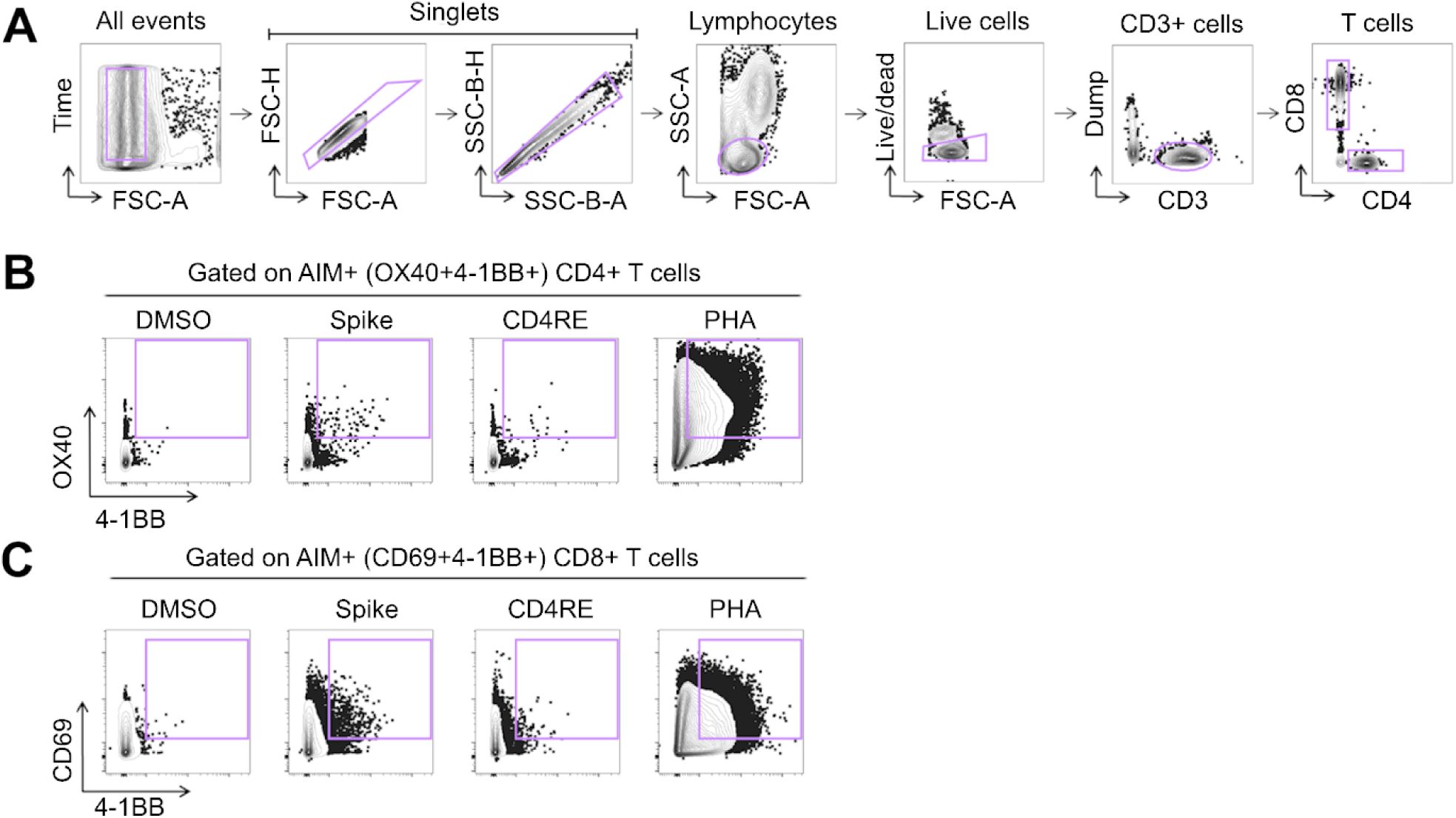
Representative gating strategy for AIM+CD4+ and CD8+ antigen-specific T cell analysis. Dot-plots show representative flow-cytometry data of gating strategies to define (**A**) bulk CD3+CD4+ and CD3+CD8+ cells or antigen-specific (**B**) AIM+ (OX40+CD137+) CD4+ T cells and (**C**) AIM+ (CD137+ CD69+) CD8+ T cells following 24h of stimulation with DMSO, Spike MP, CD4RE MP and PHA.

**Table S1.** Donor cohort demographics.

|  |  |
| --- | --- |
| Age (years) | 21 to 81 (median = 42, IQR* = 30) |
| Sex (Female/Male) | 67% (52/78) vs. 33% (26/78) |
| Race |  |
| Caucasian | 67% (52/78) |
| Hispanic or Latino | 14% (11/78) |
| Asian | 13% (10/78) |
| Black or African American | 3% (2/78) |
| Native Hawaiian or Other Pacific Islander | 1% (1/78) |
| More Than One Race | 3% (2/78) |
| Vaccine distribution | (Moderna vs. Pfizer vs. Novavax) |
| Full vaccination | 50.0% vs. 48.7% vs. 1.3% |
| 1st booster | 56.5% vs. 42.0.% vs. 1.5% |
| 2nd booster | 44.8% vs. 53.5% vs. 1.7% |

**Table S2.** Breakdown of assays performed per donor.

**Table S3.** Sample-specific metrics of Spike-reactive single-cell transcriptomes.

**Table S4.**
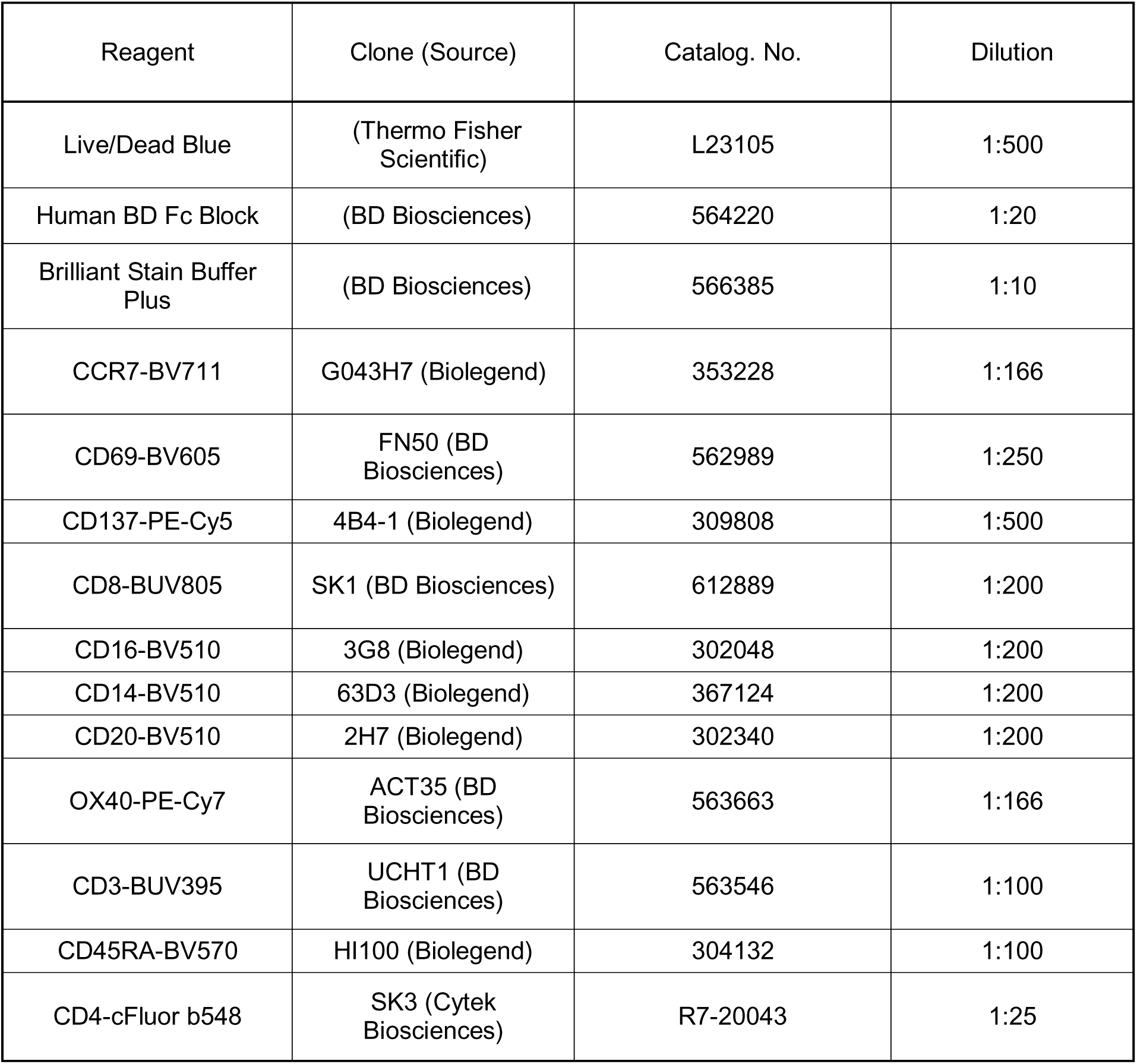
List of antibodies used by flow cytometry.

**Table S6.** Markers of Spike-reactive T cell clusters.

**Table S5.** Metrics on scRNA-seq libraries and aggregated datasets.

## REFERENCES

1. Andrews, N., Stowe, J., Kirsebom, F., Toffa, S., Rickeard, T., Gallagher, E., Gower, C., Kall, M., Groves, N., O’Connell, A.M., et al. (2022a). Covid-19 Vaccine Effectiveness against the Omicron (B.1.1.529) Variant. N Engl J Med 386, 1532–1546. 10.1056/NEJMoa2119451.

2. Andrews, N., Tessier, E., Stowe, J., Gower, C., Kirsebom, F., Simmons, R., Gallagher, E., Thelwall, S., Groves, N., Dabrera, G., et al. (2022b). Duration of Protection against Mild and Severe Disease by Covid- 19 Vaccines. N Engl J Med 386, 340–350. 10.1056/NEJMoa2115481.

3. Arlehamn, C.L., Seumois, G., Gerasimova, A., Huang, C., Fu, Z., Yue, X., Sette, A., Vijayanand, P., and Peters, B. (2014). Transcriptional profile of tuberculosis antigen-specific T cells reveals novel multifunctional features. J Immunol 193, 2931–2940. 10.4049/jimmunol.1401151.

4. Arunachalam, P.S., Lai, L., Samaha, H., Feng, Y., Hu, M., Hui, H.S., Wali, B., Ellis, M., Davis-Gardner, M.E., Huerta, C., et al. (2023). Durability of immune responses to mRNA booster vaccination against COVID-19. J Clin Invest 133. 10.1172/JCI167955.

5. Atmar, R.L., Lyke, K.E., Deming, M.E., Jackson, L.A., Branche, A.R., El Sahly, H.M., Rostad, C.A., Martin, J.M., Johnston, C., Rupp, R.E., et al. (2022). Homologous and Heterologous Covid-19 Booster Vaccinations. N Engl J Med 386, 1046–1057. 10.1056/NEJMoa2116414.

6. Azim Majumder, M.A., and Razzaque, M.S. (2022). Repeated vaccination and ’vaccine exhaustion’: relevance to the COVID-19 crisis. Expert Rev Vaccines 21, 1011–1014. 10.1080/14760584.2022.2071705.

7. Baden, L.R., El Sahly, H.M., Essink, B., Kotloff, K., Frey, S., Novak, R., Diemert, D., Spector, S.A., Rouphael, N., Creech, C.B., et al. (2021). Efficacy and Safety of the mRNA-1273 SARS-CoV-2 Vaccine. N Engl J Med 384, 403–416. 10.1056/NEJMoa2035389.

8. Bertoletti, A., Le Bert, N., and Tan, A.T. (2022). SARS-CoV-2-specific T cells in the changing landscape of the COVID-19 pandemic. Immunity 55, 1764–1778. 10.1016/j.immuni.2022.08.008.

9. Boretti, A. (2024). mRNA vaccine boosters and impaired immune system response in immune compromised individuals: a narrative review. Clin Exp Med 24, 23. 10.1007/s10238-023-01264-1.

10. Cai, C., Gao, Y., Adamo, S., Rivera-Ballesteros, O., Hansson, L., Osterborg, A., Bergman, P., Sandberg, J.K., Ljunggren, H.G., Bjorkstrom, N.K., et al. (2023). SARS-CoV-2 vaccination enhances the effector qualities of spike-specific T cells induced by COVID-19. Sci Immunol 8, eadh0687. 10.1126/sciimmunol.adh0687.

11. Chemaitelly, H., Ayoub, H.H., Tang, P., Coyle, P., Yassine, H.M., Al Thani, A.A., Al-Khatib, H.A., Hasan, M.R., Al-Kanaani, Z., Al-Kuwari, E., et al. (2023). Long-term COVID-19 booster effectiveness by infection history and clinical vulnerability and immune imprinting: a retrospective population-based cohort study. Lancet Infect Dis 23, 816–827. 10.1016/S1473-3099(23)00058-0.

12. Cohen, K.W., Linderman, S.L., Moodie, Z., Czartoski, J., Lai, L., Mantus, G., Norwood, C., Nyhoff, L.E., Edara, V.V., Floyd, K., et al. (2021). Longitudinal analysis shows durable and broad immune memory after SARS-CoV-2 infection with persisting antibody responses and memory B and T cells. Cell Rep Med 2, 100354. 10.1016/j.xcrm.2021.100354.

13. da Silva Antunes, R., Weiskopf, D., Sidney, J., Rubiro, P., Peters, B., Lindestam Arlehamn, C.S., Grifoni, A., and Sette, A. (2023). The MegaPool Approach to Characterize Adaptive CD4+ and CD8+ T Cell Responses. Curr Protoc 3, e934. 10.1002/cpz1.934.

14. Daamen, A.R., Bachali, P., Bonham, C.A., Somerville, L., Sturek, J.M., Grammer, A.C., Kadl, A., and Lipsky, P.E. (2022). COVID-19 patients exhibit unique transcriptional signatures indicative of disease severity. Front Immunol 13, 989556. 10.3389/fimmu.2022.989556.

15. Dan, J.M., Lindestam Arlehamn, C.S., Weiskopf, D., da Silva Antunes, R., Havenar-Daughton, C., Reiss, S.M., Brigger, M., Bothwell, M., Sette, A., and Crotty, S. (2016). A Cytokine-Independent Approach To Identify Antigen-Specific Human Germinal Center T Follicular Helper Cells and Rare Antigen-Specific CD4+ T Cells in Blood. J Immunol 197, 983–993. 10.4049/jimmunol.1600318.

16. Dan, J.M., Mateus, J., Kato, Y., Hastie, K.M., Yu, E.D., Faliti, C.E., Grifoni, A., Ramirez, S.I., Haupt, S., Frazier, A., et al. (2021a). Immunological memory to SARS-CoV-2 assessed for up to 8 months after infection. Science 371. 10.1126/science.abf4063.

17. Dan, J.M., Mateus, J., Kato, Y., Hastie, K.M., Yu, E.D., Faliti, C.E., Grifoni, A., Ramirez, S.I., Haupt, S., Frazier, A., et al. (2021b). Immunological memory to SARS-CoV-2 assessed for up to 8 months after infection. Science 371, eabf4063. doi:10.1126/science.abf4063.

18. Davis-Porada, J., George, A.B., Lam, N., Caron, D.P., Gray, J.I., Huang, J., Hwu, J., Wells, S.B., Matsumoto, R., Kubota, M., et al. (2024). Maintenance and functional regulation of immune memory to COVID-19 vaccines in tissues. Immunity. 10.1016/j.immuni.2024.10.003.

19. Eschweiler, S., Clarke, J., Ramirez-Suastegui, C., Panwar, B., Madrigal, A., Chee, S.J., Karydis, I., Woo, E., Alzetani, A., Elsheikh, S., et al. (2021). Intratumoral follicular regulatory T cells curtail anti-PD-1 treatment efficacy. Nat Immunol 22, 1052–1063. 10.1038/s41590-021-00958-6.

20. Finak, G., McDavid, A., Yajima, M., Deng, J., Gersuk, V., Shalek, A.K., Slichter, C.K., Miller, H.W., McElrath, M.J., Prlic, M., et al. (2015). MAST: a flexible statistical framework for assessing transcriptional changes and characterizing heterogeneity in single-cell RNA sequencing data. Genome Biol 16, 278. 10.1186/s13059-015-0844-5.

21. Franco, A., Song, J., Chambers, C., Sette, A., and Grifoni, A. (2023). SARS-CoV-2 spike-specific regulatory T cells (Treg) expand and develop memory in vaccine recipients suggesting a role for immune regulation in preventing severe symptoms in COVID-19. Autoimmunity 56, 2259133. 10.1080/08916934.2023.2259133.

22. Galbraith, M.D., Kinning, K.T., Sullivan, K.D., Araya, P., Smith, K.P., Granrath, R.E., Shaw, J.R., Baxter, R., Jordan, K.R., Russell, S., et al. (2022). Specialized interferon action in COVID-19. Proc Natl Acad Sci U S A 119. 10.1073/pnas.2116730119.

23. Geers, D., Shamier, M.C., Bogers, S., den Hartog, G., Gommers, L., Nieuwkoop, N.N., Schmitz, K.S., Rijsbergen, L.C., van Osch, J.A.T., Dijkhuizen, E., et al. (2021). SARS-CoV-2 variants of concern partially escape humoral but not T-cell responses in COVID-19 convalescent donors and vaccinees. Sci Immunol 6. 10.1126/sciimmunol.abj1750.

24. Gray-Gaillard, S.L., Solis, S.M., Chen, H.M., Monteiro, C., Ciabattoni, G., Samanovic, M.I., Cornelius, A.R., Williams, T., Geesey, E., Rodriguez, M., et al. (2024). SARS-CoV-2 inflammation durably imprints memory CD4 T cells. Sci Immunol 9, eadj8526. 10.1126/sciimmunol.adj8526.

25. Grifoni, A., Weiskopf, D., Ramirez, S.I., Mateus, J., Dan, J.M., Moderbacher, C.R., Rawlings, S.A., Sutherland, A., Premkumar, L., Jadi, R.S., et al. (2020a). Targets of T Cell Responses to SARS-CoV-2 Coronavirus in Humans with COVID-19 Disease and Unexposed Individuals. Cell 181, 1489–1501.e1415. 10.1016/j.cell.2020.05.015.

26. Grifoni, A., Weiskopf, D., Ramirez, S.I., Mateus, J., Dan, J.M., Moderbacher, C.R., Rawlings, S.A., Sutherland, A., Premkumar, L., Jadi, R.S., et al. (2020b). Targets of T Cell Responses to SARS-CoV-2 Coronavirus in Humans with COVID-19 Disease and Unexposed Individuals. Cell 181, 1489–1501 e1415. 10.1016/j.cell.2020.05.015.

27. Jo, N., Hidaka, Y., Kikuchi, O., Fukahori, M., Sawada, T., Aoki, M., Yamamoto, M., Nagao, M., Morita, S., Nakajima, T.E., et al. (2023). Impaired CD4(+) T cell response in older adults is associated with reduced immunogenicity and reactogenicity of mRNA COVID-19 vaccination. Nat Aging 3, 82–92. 10.1038/s43587-022-00343-4.

28. Jovanovic, M., Sekulic, S., Jocic, M., Jurisevic, M., Gajovic, N., Jovanovic, M., Arsenijevic, N., Jovanovic, M., Mijailovic, M., Milosavljevic, M., and Jovanovic, I. (2023). Increased Pro Th1 And Th17 Transcriptional Activity In Patients With Severe COVID-19. Int J Med Sci 20, 530–541. 10.7150/ijms.80498.

29. Kedzierska, K., and Thomas, P.G. (2022). Count on us: T cells in SARS-CoV-2 infection and vaccination. Cell Rep Med 3, 100562. 10.1016/j.xcrm.2022.100562.

30. Keeton, R., Tincho, M.B., Suzuki, A., Benede, N., Ngomti, A., Baguma, R., Chauke, M.V., Mennen, M., Skelem, S., Adriaanse, M., et al. (2023). Impact of SARS-CoV-2 exposure history on the T cell and IgG response. Cell Rep Med 4, 100898. 10.1016/j.xcrm.2022.100898.

31. Kim, T.S., and Shin, E.C. (2019). The activation of bystander CD8(+) T cells and their roles in viral infection. Exp Mol Med 51, 1–9. 10.1038/s12276-019-0316-1.

32. Korotkevich, G., Sukhov, V., Budin, N., Shpak, B., Artyomov, M.N., and Sergushichev, A. (2021). Fast gene set enrichment analysis. BioRxiv.

33. Koutsakos, M., Reynaldi, A., Lee, W.S., Nguyen, J., Amarasena, T., Taiaroa, G., Kinsella, P., Liew, K.C., Tran, T., Kent, H.E., et al. (2023). SARS-CoV-2 breakthrough infection induces rapid memory and de novo T cell responses. Immunity 56, 879–892 e874. 10.1016/j.immuni.2023.02.017.

34. Kusnadi, A., Ramirez-Suastegui, C., Fajardo, V., Chee, S.J., Meckiff, B.J., Simon, H., Pelosi, E., Seumois, G., Ay, F., Vijayanand, P., and Ottensmeier, C.H. (2021). Severely ill COVID-19 patients display impaired exhaustion features in SARS-CoV-2-reactive CD8(+) T cells. Sci Immunol 6. 10.1126/sciimmunol.abe4782.

35. Le Bert, N., and Samandari, T. (2024). Silent battles: immune responses in asymptomatic SARS-CoV-2 infection. Cell Mol Immunol 21, 159–170. 10.1038/s41423-024-01127-z.

36. Levin, E.G., Lustig, Y., Cohen, C., Fluss, R., Indenbaum, V., Amit, S., Doolman, R., Asraf, K., Mendelson, E., Ziv, A., et al. (2021). Waning Immune Humoral Response to BNT162b2 Covid-19 Vaccine over 6 Months. N Engl J Med 385, e84. 10.1056/NEJMoa2114583.

37. Machkovech, H.M., Hahn, A.M., Garonzik Wang, J., Grubaugh, N.D., Halfmann, P.J., Johnson, M.C., Lemieux, J.E., O’Connor, D.H., Piantadosi, A., Wei, W., and Friedrich, T.C. (2024). Persistent SARS-CoV-2 infection: significance and implications. Lancet Infect Dis 24, e453–e462. 10.1016/S1473-3099(23)00815-0.

38. Meckiff, B.J., Ramirez-Suastegui, C., Fajardo, V., Chee, S.J., Kusnadi, A., Simon, H., Eschweiler, S., Grifoni, A., Pelosi, E., Weiskopf, D., et al. (2020). Imbalance of Regulatory and Cytotoxic SARS-CoV-2-Reactive CD4(+) T Cells in COVID-19. Cell 183, 1340–1353 e1316. 10.1016/j.cell.2020.10.001.

39. Moss, P. (2022). The T cell immune response against SARS-CoV-2. Nat Immunol 23, 186–193. 10.1038/s41590-021-01122-w.

40. Mudd, P.A., Minervina, A.A., Pogorelyy, M.V., Turner, J.S., Kim, W., Kalaidina, E., Petersen, J., Schmitz, A.J., Lei, T., Haile, A., et al. (2022). SARS-CoV-2 mRNA vaccination elicits a robust and persistent T follicular helper cell response in humans. Cell 185, 603–613 e615. 10.1016/j.cell.2021.12.026.

41. Painter, M.M., Johnston, T.S., Lundgreen, K.A., Santos, J.J.S., Qin, J.S., Goel, R.R., Apostolidis, S.A., Mathew, D., Fulmer, B., Williams, J.C., et al. (2023). Prior vaccination enhances immune responses during SARS-CoV-2 breakthrough infection with early activation of memory T cells followed by production of potent neutralizing antibodies. bioRxiv. 10.1101/2023.02.05.527215.

42. Polack, F.P., Thomas, S.J., Kitchin, N., Absalon, J., Gurtman, A., Lockhart, S., Perez, J.L., Perez Marc, G., Moreira, E.D., Zerbini, C., et al. (2020). Safety and Efficacy of the BNT162b2 mRNA Covid-19 Vaccine. N Engl J Med 383, 2603–2615. 10.1056/NEJMoa2034577.

43. Pusnik, J., Monzon-Posadas, W.O., Zorn, J., Peters, K., Baum, M., Proksch, H., Schluter, C.B., Alter, G., Menting, T., and Streeck, H. (2023). SARS-CoV-2 humoral and cellular immunity following different combinations of vaccination and breakthrough infection. Nat Commun 14, 572. 10.1038/s41467-023- 36250-4.

44. Ramirez, S.I., Faraji, F., Hills, L.B., Lopez, P.G., Goodwin, B., Stacey, H.D., Sutton, H.J., Hastie, K.M., Saphire, E.O., Kim, H.J., et al. (2024). Immunological memory diversity in the human upper airway. Nature 632, 630–636. 10.1038/s41586-024-07748-8.

45. Reinscheid, M., Luxenburger, H., Karl, V., Graeser, A., Giese, S., Ciminski, K., Reeg, D.B., Oberhardt, V., Roehlen, N., Lang-Meli, J., et al. (2022). COVID-19 mRNA booster vaccine induces transient CD8+ T effector cell responses while conserving the memory pool for subsequent reactivation. Nat Commun 13, 4631. 10.1038/s41467-022-32324-x.

46. Riou, C., Keeton, R., Moyo-Gwete, T., Hermanus, T., Kgagudi, P., Baguma, R., Valley-Omar, Z., Smith, M., Tegally, H., Doolabh, D., et al. (2022). Escape from recognition of SARS-CoV-2 variant spike epitopes but overall preservation of T cell immunity. Sci Transl Med 14, eabj6824. 10.1126/scitranslmed.abj6824.

47. Schneider, K., Loewendorf, A., De Trez, C., Fulton, J., Rhode, A., Shumway, H., Ha, S., Patterson, G., Pfeffer, K., Nedospasov, S.A., et al. (2008). Lymphotoxin-mediated crosstalk between B cells and splenic stroma promotes the initial type I interferon response to cytomegalovirus. Cell Host Microbe 3, 67–76. 10.1016/j.chom.2007.12.008.

48. Schneider, W.M., Chevillotte, M.D., and Rice, C.M. (2014). Interferon-stimulated genes: a complex web of host defenses. Annu Rev Immunol 32, 513–545. 10.1146/annurev-immunol-032713-120231.

49. Sette, A., and Crotty, S. (2022). Immunological memory to SARS-CoV-2 infection and COVID-19 vaccines. Immunol Rev 310, 27–46. 10.1111/imr.13089.

50. Sette, A., Sidney, J., and Crotty, S. (2023). T Cell Responses to SARS-CoV-2. Annu Rev Immunol 41, 343–373. 10.1146/annurev-immunol-101721-061120.

51. Shaikh, N., Swali, P., and Houben, R. (2023). Asymptomatic but infectious - The silent driver of pathogen transmission. A pragmatic review. Epidemics 44, 100704. 10.1016/j.epidem.2023.100704.

52. Srivastava, K., Carreno, J.M., Gleason, C., Monahan, B., Singh, G., Abbad, A., Tcheou, J., Raskin, A., Kleiner, G., van Bakel, H., et al. (2024). SARS-CoV-2-infection- and vaccine-induced antibody responses are long lasting with an initial waning phase followed by a stabilization phase. Immunity 57, 587–599 e584. 10.1016/j.immuni.2024.01.017.

53. Stuart, T., Butler, A., Hoffman, P., Hafemeister, C., Papalexi, E., Mauck, W.M., 3rd, Hao, Y., Stoeckius, M., Smibert, P., and Satija, R. (2019). Comprehensive Integration of Single-Cell Data. Cell 177, 1888–1902 e1821. 10.1016/j.cell.2019.05.031.

54. Suresh, M., Lanier, G., Large, M.K., Whitmire, J.K., Altman, J.D., Ruddle, N.H., and Ahmed, R. (2002). Role of lymphotoxin alpha in T-cell responses during an acute viral infection. J Virol 76, 3943–3951. 10.1128/jvi.76.8.3943-3951.2002.

55. Tan, A.T., Linster, M., Tan, C.W., Le Bert, N., Chia, W.N., Kunasegaran, K., Zhuang, Y., Tham, C.Y.L., Chia, A., Smith, G.J.D., et al. (2021). Early induction of functional SARS-CoV-2-specific T cells associates with rapid viral clearance and mild disease in COVID-19 patients. Cell Rep 34, 108728. 10.1016/j.celrep.2021.108728.

56. Tarke, A., Potesta, M., Varchetta, S., Fenoglio, D., Iannetta, M., Sarmati, L., Mele, D., Dentone, C., Bassetti, M., Montesano, C., et al. (2022). Early and Polyantigenic CD4 T Cell Responses Correlate with Mild Disease in Acute COVID-19 Donors. Int J Mol Sci 23. 10.3390/ijms23137155.

57. Tarke, A., Ramezani-Rad, P., Alves Pereira Neto, T., Lee, Y., Silva-Moraes, V., Goodwin, B., Bloom, N., Siddiqui, L., Avalos, L., Frazier, A., et al. (2024). SARS-CoV-2 breakthrough infections enhance T cell response magnitude, breadth, and epitope repertoire. Cell Rep Med 5, 101583. 10.1016/j.xcrm.2024.101583.

58. Tarke, A., Sidney, J., Methot, N., Yu, E.D., Zhang, Y., Dan, J.M., Goodwin, B., Rubiro, P., Sutherland, A., Wang, E., et al. (2021). Impact of SARS-CoV-2 variants on the total CD4(+) and CD8(+) T cell reactivity in infected or vaccinated individuals. Cell Rep Med 2, 100355. 10.1016/j.xcrm.2021.100355.

59. Wang, B., Andraweera, P., Elliott, S., Mohammed, H., Lassi, Z., Twigger, A., Borgas, C., Gunasekera, S., Ladhani, S., and Marshall, H.S. (2023). Asymptomatic SARS-CoV-2 Infection by Age: A Global Systematic Review and Meta-analysis. Pediatr Infect Dis J 42, 232–239. 10.1097/INF.0000000000003791.

60. Yu, E.D., Narowski, T.M., Wang, E., Garrigan, E., Mateus, J., Frazier, A., Weiskopf, D., Grifoni, A., Premkumar, L., da Silva Antunes, R., and Sette, A. (2022a). Immunological memory to common cold coronaviruses assessed longitudinally over a three-year period pre-COVID19 pandemic. Cell Host Microbe 30, 1269–1278 e1264. 10.1016/j.chom.2022.07.012.

61. Yu, E.D., Wang, E., Garrigan, E., Goodwin, B., Sutherland, A., Tarke, A., Chang, J., Galvez, R.I., Mateus, J., Ramirez, S.I., et al. (2022b). Development of a T cell-based immunodiagnostic system to effectively distinguish SARS-CoV-2 infection and COVID-19 vaccination status. Cell Host Microbe 30, 388–399 e383. 10.1016/j.chom.2022.02.003.

62. Yu, W., Guo, Y., Zhang, S., Kong, Y., Shen, Z., and Zhang, J. (2022c). Proportion of asymptomatic infection and nonsevere disease caused by SARS-CoV-2 Omicron variant: A systematic review and analysis. J Med Virol 94, 5790–5801. 10.1002/jmv.28066.

63. Zhang, B., Upadhyay, R., Hao, Y., Samanovic, M.I., Herati, R.S., Blair, J.D., Axelrad, J., Mulligan, M.J., Littman, D.R., and Satija, R. (2023). Multimodal single-cell datasets characterize antigen-specific CD8(+) T cells across SARS-CoV-2 vaccination and infection. Nat Immunol 24, 1725–1734. 10.1038/s41590-023-01608-9.

64. Zhang, Z., Mateus, J., Coelho, C.H., Dan, J.M., Moderbacher, C.R., Galvez, R.I., Cortes, F.H., Grifoni, A., Tarke, A., Chang, J., et al. (2022a). Humoral and cellular immune memory to four COVID-19 vaccines. Cell 185, 2434–2451 e2417. 10.1016/j.cell.2022.05.022.

65. Zhang, Z., Mateus, J., Coelho, C.H., Dan, J.M., Moderbacher, C.R., Gálvez, R.I., Cortes, F.H., Grifoni, A., Tarke, A., Chang, J., et al. (2022b). Humoral and cellular immune memory to four COVID-19 vaccines. Cell. 10.1016/j.cell.2022.05.022.

